# ALDH1B1 promotes mitochondrial γ-butyrobetaine biosynthesis and fatty acid oxidation in colorectal cancer

**DOI:** 10.64898/2026.09.24.753323

**Authors:** Zhiping Feng, Thomas E. Bearrood, Sydney L. Campbell, Adrianne M. Kinsey, Nedas Matulionis, Che-Hong Chen, Takuya Seike, Logan Leak, A. Kaan Tarhan, Scott J. Dixon, Daria Mochly-Rosen, Heather R. Christofk, James K. Chen

## Abstract

Metabolic reprogramming is a hallmark of cancer cells, and stem-like populations often upregulate aldehyde dehydrogenases (ALDHs). Functional studies have established essential roles for individual ALDH isoforms in tumor initiation, progression, and metastasis; however, the mechanisms by which these enzymes promote malignancy remain poorly understood. Here we show that aldehyde dehydrogenase 1B1 (ALDH1B1), a mitochondrial enzyme highly expressed in colorectal cancer (CRC) and pancreatic ductal adenocarcinoma (PDAC), generates γ-butyrobetaine (GBB) and γ-aminobutyric acid (GABA) in CRC cells. The biosynthesis of both aminocarboxylic acids has been attributed to the cytosolic enzyme ALDH9A1, and we demonstrate that mitochondrial GBB and GABA are functionally distinct from the cytosolic pools of these metabolites. We further demonstrate that mitochondrial GBB can function as an antiport substrate of carnitine-acylcarnitine translocase (CACT), the transporter that mediates fatty acid uptake into the mitochondrial inner matrix. This activity complements the role of cytosolic GBB as the biosynthetic precursor to carnitine. Accordingly, ALDH1B1 can markedly enhance mitochondrial fatty acid oxidation (FAO), a catabolic process that has been linked to CRC and PDAC stemness, progression, and metastasis. Our findings reveal an unexpected role for ALDH1B1 in carnitine metabolism, GABA biosynthesis, and FAO and illustrate how the compartmental reprogramming of metabolic pathways can promote tumor growth.

## INTRODUCTION

Cancer cells can undergo extensive metabolic adaptations to enable their survival, proliferation, and resistance to therapy. Early studies of cancer metabolism centered on the Warburg effect, whereby tumor cells maintain high rates of glycolysis and lactate production despite oxygen availability ^1,2^. While this phenomenon remains a defining feature of many cancers, it is now recognized that tumor cells possess remarkable bioenergetic plasticity, enabling them to exploit diverse nutrient sources and adapt to hostile microenvironments such as hypoxia, severe oxidative stress, and nutrient deprivation^3^. This metabolic flexibility is not uniformly distributed across tumor cells, and distinct cell subpopulations employ specialized strategies to support specific functional states^4,5^.

A critical axis of this metabolic heterogeneity involves rare subpopulations of tumor cells that exhibit self-renewal capacity, tumor-initiating potential, and resistance to conventional therapies^6,7^. Commonly referred to as cancer stem cells (CSCs), these slow-cycling tumor cells have bioenergetic states that are distinct from those of differentiated cell lineages. Aldehyde dehydrogenases (ALDHs) are uniquely positioned at this nexus of metabolism and stemness. This superfamily of NAD(P)⁺-dependent enzymes comprises 19 human isoforms with diverse substrate specificities, tissue expression patterns, and subcellular localizations, and elevated ALDH activity is one of the most widely used markers for isolating and identifying stem cells in normal and malignant tissues^8^. In particular, members of the ALDH1 subfamily have been associated with the maintenance of normal stem cell populations in tissues spanning hematopoietic lineages, neural stem cells, and mammary epithelium^9^. High ALDH1 activity also correlates with tumor resistance to chemotherapy and radiation, metastasis, and poor patient prognoses^10,11^, and individual ALDH1 enzymes have been linked to specific cancer types. Our mechanistic understanding of oncogenic ALDH1 functions has largely been derived from studies of the cytosolic isoforms ALDH1A1 and ALDH1A3. For example, ALDH1A1 can drive angiogenesis in breast cancer through retinoic acid biosynthesis and nuclear hormone receptor-dependent activation of HIF-1α/VEGF^12^. ALDH1A3 similarly drives breast cancer growth and metastasis through retinoic acid signaling^13^, and in melanoma, it forms a nuclear enzymatic complex with acetyl-CoA synthetase 2 to reprogram glucose metabolism and modulate histone H3 acetylation^14^.

Mitochondrial ALDH1 isoforms have also emerged as important contributors to cancer biology, paralleling growing evidence that mitochondrial metabolism supports CSC maintenance and function^15–17^. For instance, ALDH1B1 is highly expressed in colorectal cancer (CRC)^18,19^ and pancreatic ductal adenocarcinoma (PDAC)^20,21^, and the enzyme is a key metabolic dependency of these gastrointestinal malignancies. Genetic loss of *Aldh1b1* in mice markedly reduces CRC progression in adenomatous polyposis coli (*Apc*) mutant mice^22^ and entirely abrogates PDAC initiation driven by oncogenic KRAS(G12D)^21^. In addition, SW480 CRC cells exhibit a greater dependency on ALDH1B1 when cultured as three-dimensional (3D) spheroids rather than two-dimensional (2D) monolayers^23^. Since 3D cancer cell cultures have been shown to enrich for and require stem-like cells^24–27^, these results suggest that ALDH1B1 supports stemness-associated programs. Consistent with this idea, both genetic and pharmacological inhibition of ALDH1B1 in SW480 cells reduces expression of the CRC stem cell-associated markers *keratin 15* (*KRT15*) and *doublecortin-like kinase 1* (*DCLK1*)^23,28,29^, and ALDH1B1 is upregulated in normal stem and progenitor cell populations of the intestine and pancreas^21,30,31^.

While these findings establish a key role for ALDH1B1 in CRC and PDAC, the precise molecular and cellular functions of ALDH1B1 remain enigmatic. Knockdown of *ALDH1B1* reduces Wnt, Notch, and PI3K/Akt signaling in CRC cells^31^, but the mechanisms that link ALDH1B1 enzymatic activity to these pathways is unknown. A recent report also proposes that ALDH1B1 regulates ferroptosis in sarcoma cells by oxidizing 4-hydroxynonenal (4-HNE)^32^, but whether 4-HNE constitutes a bona fide physiological substrate for ALDH1B1 remains unclear.

In this work, we have coupled chemical ALDH1B1 inhibition with metabolomics to identify γ-butyrobetaine (GBB) and γ-aminobutyric acid (GABA) as metabolic products of ALDH1B1 in CRC cells. GBB and GABA biosynthesis have previously been attributed to cytosolic enzyme ALDH9A1, and our studies indicate that ALDH1B1 generates mitochondrial pools of these metabolites that are functionally distinct. In the case of GBB, the cytosolic metabolite is a critical intermediate in the biosynthesis of L-carnitine, which serves as the obligate carrier for importing long-chain fatty acids into mitochondria via carnitine-acylcarnitine translocase (CACT). The fatty acids then undergo β-oxidation to yield NADH, FADH_2_, and acetyl-CoA. Rather than contributing directly to carnitine biosynthesis, mitochondrial GBB can first serve as an antiport substrate for CACT, increasing fatty acid transport and oxidation. Accordingly, we observe that ALDH1B1 can markedly enhances fatty acid oxidation (FAO) activity in CRC spheroids. Taken together, our studies uncover a role for ALDH1B1 in the subcellular reprogramming of GBB biosynthesis, thereby shifting CRC cells to an FAO-dependent state that promotes stemness and tumor growth.

## RESULTS

### ALDH1B1 does not regulate ferroptosis in CRC cells

We began our studies by assessing previously reported functions of ALDH1B1 in cancer cells. A recent study concluded that ALDH1B1 regulates ferroptosis in HT-1080 fibrosarcoma cells by oxidizing 4-HNE; transduction of this epithelial line with *ALDH1B1*-targeting short hairpin RNAs (shRNAs) or treatment with the pan-ALDH inhibitor *N*,*N*-diethylaminobenzaldehyde (DEAB) sensitized the cells to lipid peroxidation-driven cell death^32^. We aimed to recapitulate these findings using both DEAB and IGUANA-5, a highly specific ALDH1B1 inhibitor that we recently developed through structure-guided medicinal chemistry (Fig. 1A and Supplementary Fig. 1; biochemical IC_50_ = 13 nM)^33^. Unexpectedly, treating HT-1080 cells with either compound did not increase their sensitivity to RSL3, a glutathione peroxidase 4 inhibitor and ferroptosis inducer^34^, as assessed by ATP- and SYTOX Green-based cell viability assays (Fig. 1B and Supplementary Fig. 2A-B).

**Figure 1.**
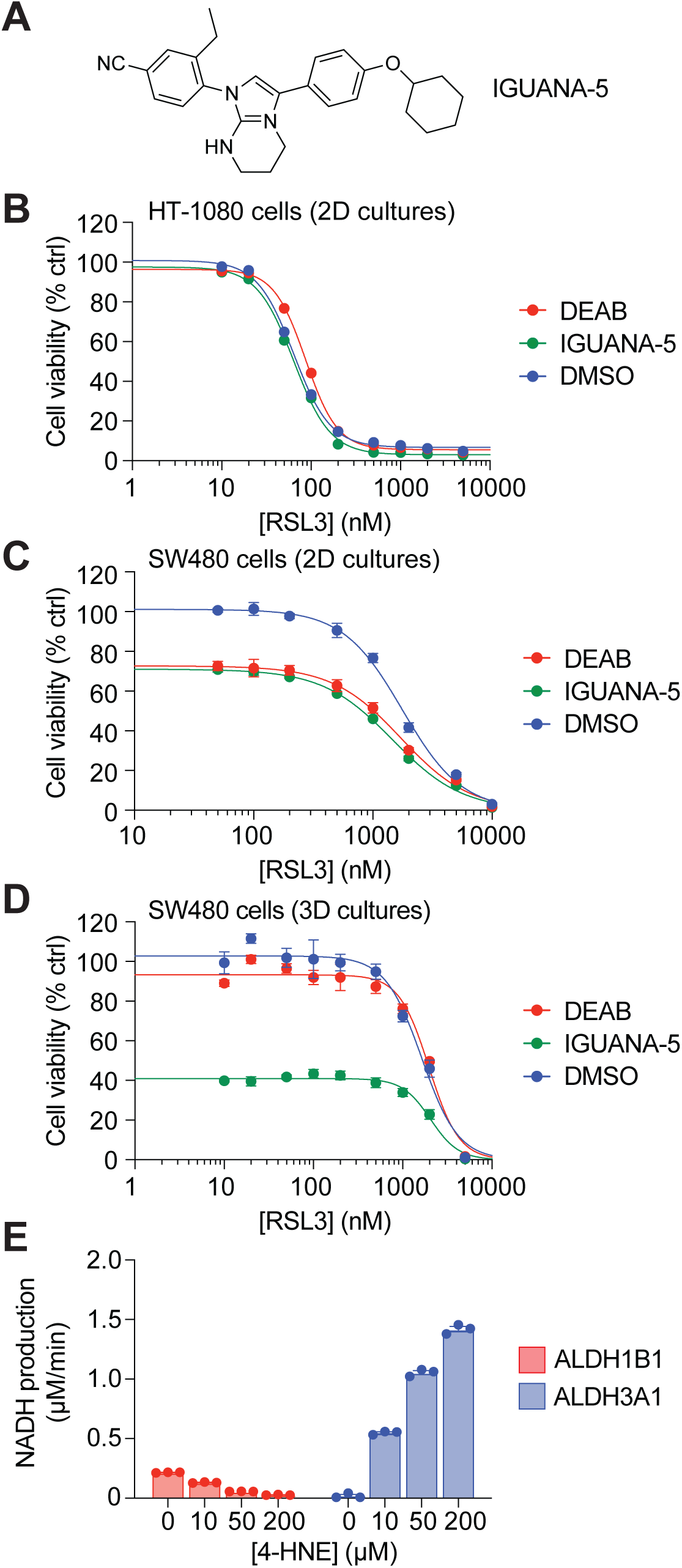
ALDH1B1 does not regulate ferroptosis in CRC cells. (A) Chemical structure of IGUANA-5, a potent and selective ALDH1B1 inhibitor. (B-D) Dose-response curves for the ferroptosis inducer RSL3 in HT-1080 cells (B) and SW480 cells (C-D) in the indicated culture conditions and co-treated with DMSO, 100 µM DEAB or 2 µM IGUANA-5. Adherent and spheroid cultures were treated for 72 h and 4 days, respectively. IC_50_ values for the DMSO, DEAB, and IGUANA-5 treatment conditions were determined to be 64, 87, and 64 nM in (B); 1.8, 1.7, and 1.5 µM in (C); and 1.9, 2.3, and 2.3 µM in (D). Cell viability measurements were normalized to the DMSO-treated group. (E) Enzymatic activities of ALDH1B1 (100 nM monomer concentration) and ALDH3A1 (5 nM monomer concentration) against 4-HNE. All graphed data are the average of three biological replicates ± s.e.m.

We then investigated whether ALDH1B1 influences ferroptosis sensitivity in CRC cells. Mirroring our results with the fibrosarcoma line, neither IGUANA-5 nor DEAB made SW480 cells more susceptible to RSL3-induced ferroptosis (Fig. 1C-D). We also examined the potential role of ALDH1B1 in suppressing CRC cell ferroptosis by treating *ALDH1B1^−/−^* SW480 spheroids with the ferroptosis inhibitors, ferrostatin-1^35^ and liproxstatin-1^36^. Both compounds failed to rescue the viability of these 3D cultures, suggesting ferroptosis does not contribute to this *ALDH1B1* knockout phenotype (Supplementary Fig. 2C-D). These results prompted us to evaluate the ability of ALDH1B1 to utilize 4-HNE as a substrate. We incubated recombinant human ALDH1B1 with 4-HNE in the presence of NAD^+^, measuring NADH fluorescence to track 4-HNE oxidation and using human ALDH3A1 as a positive control^37^. ALDH3A1, but not ALDH1B1, efficiently oxidized this lipid peroxidation product. In fact, 4-HNE appeared to inhibit background ALDH1B1 activity in a concentration-dependent manner (Fig. 1E).

ALDH1B1 has also been proposed to oxidize all-*trans* retinal as a means of modulating retinoic acid signaling-dependent differentiation of hematopoietic stem cells^38,39^. We therefore used a diaphorase-coupled assay to monitor this potential enzymatic reaction (all-*trans* retinal and NADH have overlapping absorbance spectra). We found all-*trans* retinal to be a very poor substrate for ALDH1B1 in comparison to ALDH1A3, a well-established retinal dehydrogenase^40^ (Supplementary Fig. 3).

### Identification of ALDH1B1 substrates

To re-examine the physiological roles of ALDH1B1, we interrogated the ALDH1B1-dependent metabolome in SW480 cells. For these studies, we leveraged our ALDH1B1 isoform-specific inhibitor IGUANA-5, exploiting the temporal control afforded by small molecules and minimizing the compensatory adaptations and other secondary effects that often accompany genetic perturbations. To discern on-target versus off-target activities of the inhibitor, we cultured *ALDH1B1^−/−^* SW480 cells transduced with either exogenous *ALDH1B1* or *EGFP*^23^ and treated the lines with either 2 µM IGUANA-5 or DMSO vehicle alone for 12 h. We anticipated that ALDH1B1-dependent changes induced by IGUANA-5 treatment would only be observed in the *ALDH1B1^−/−^* SW480 cells rescued with exogenous ALDH1B1 but not in those overexpressing EGFP. We used LC-MS/MS to profile 157 metabolites spanning diverse biochemical pathways (Supplementary File 1), observing that IGUANA-5 treatment downregulated several trimethylammonium-containing metabolites in the *ALDH1B1*-transduced knockout line, including GBB, carnitine, acetylcarnitine, propionylcarnitine, and betaine (Fig. 2A). The ALDH1B1 inhibitor also selectively decreased levels of the structurally similar metabolite GABA and upregulated nucleoside levels in these cells. In comparison, only the nucleosides were significantly affected by IGUANA-5 in EGFP-expressing *ALDH1B1^−/−^* cells (Fig. 2B), indicating that this metabolic change is an off-target effect of this small molecule.

**Figure 2.**
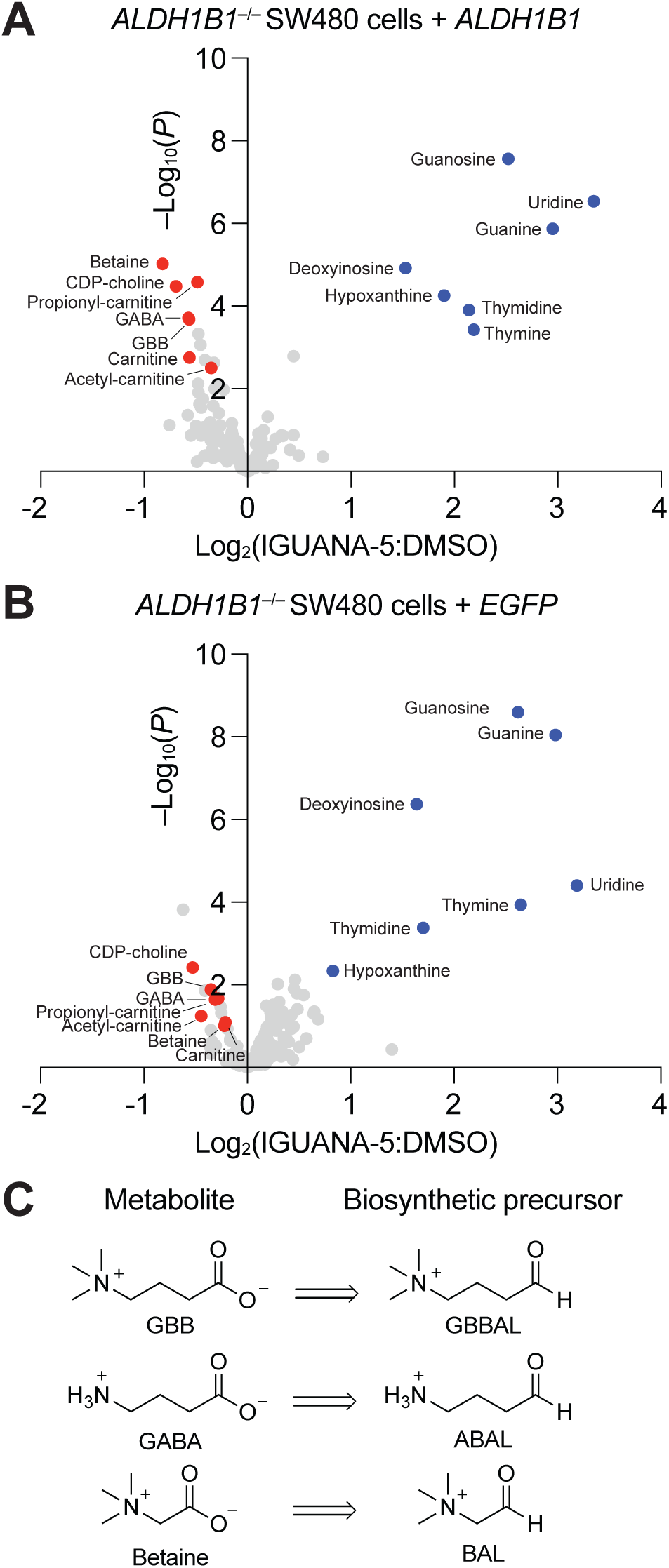
Profiling of ALDH1B1-dependent metabolites in SW480 cells. (A-B) Metabolic changes in *ALDH1B1^−/−^* SW480 cells overexpressing exogenous ALDH1B1 (A) or EGFP (B) upon treatment with 2 µM IGUANA-5 or DMSO for 12 h. Each dot represents an individual metabolite, indicating the ratio of the normalized mass spectrometry peak areas for the IGUANA-5-versus DMSO-treated samples. Data are the average of five biological replicates. On-target effects of IGUANA-5 on the levels of trimethylammonium-containing metabolites and GABA are shown in red, and off-target effects on nucleosides in blue. (C) Chemical structures of carboxylic acid metabolites identified in the profiling study and their aldehyde precursors.

Among the metabolites with ALDH1B1 dependency, GBB, betaine, and GABA are carboxylic acids produced through oxidation of their corresponding aldehydes: γ-butyrobetaine aldehyde (GBBAL), betaine aldehyde (BAL), and γ-aminobutyraldehyde (also known as 4-aminobutanal; ABAL), respectively (Fig. 2C). GBB and GABA biosynthesis have been attributed to cytosolic ALDH9A1, and betaine production has been associated with both ALDH9A1 and ALDH7A1, an isoform that localizes to multiple cellular compartments^41–45^. Since our metabolite profiling study did not include aldehydes due to their high reactivity, we assessed whether ALDH1B1 can also oxidize the aminoaldehyde precursors for GBB, GABA, and betaine. We first synthesized GBBAL and evaluated it as an enzymatic substrate for bacterially expressed human ALDH1B1 and ALDH9A1. ALDH1B1 efficiently oxidized GBBAL with a *K*_m_ value comparable to that of ALDH9A1, but ALDH1B1 had a substantially higher turnover rate (*k_cat_*) (Fig. 3A). We further evaluated the extent to which other ALDH isoforms can accept GBBAL as a substrate. Among the ten paralogs tested, only ALDH1B1 and ALDH9A1 efficiently oxidized GBBAL, and ALDH1A1 displayed minimal activity that increased with GBBAL concentration (Fig. 3B). We similarly observed that ABAL is readily oxidized by ALDH1B1, with a *k_cat_*/*K_m_* ratio similar to that of ALDH1B1/GBBAL and over 10-fold greater than that of ALDH9A1/ABAL (Fig. 3C). ALDH1B1 was also the most efficient dehydrogenase for ABAL among the ten ALDH family members we evaluated, with ALDH1A1, ALDH2, and ALDH9A1 accommodating this aldehyde substrate to a limited extent (Fig. 3D). However, ALDH1B1 oxidized BAL with a much lower *k_cat_*/*K_m_* ratio, indicating that this short-chain trimethylammonium aldehyde is a relatively poor substrate for this ALDH family member (Fig. 3E). In comparison, ALDH9A1 efficiently converted BAL into betaine, and we unexpectedly found that ALDH7A1 was inactive against BAL even though it could oxidize acetaldehyde (Supplementary Fig. 4). Taken together, these findings establish ALDH1B1/GBBAL and ALDH1B1/ABAL as preferred enzyme/substrate pairs.

**Figure 3.**
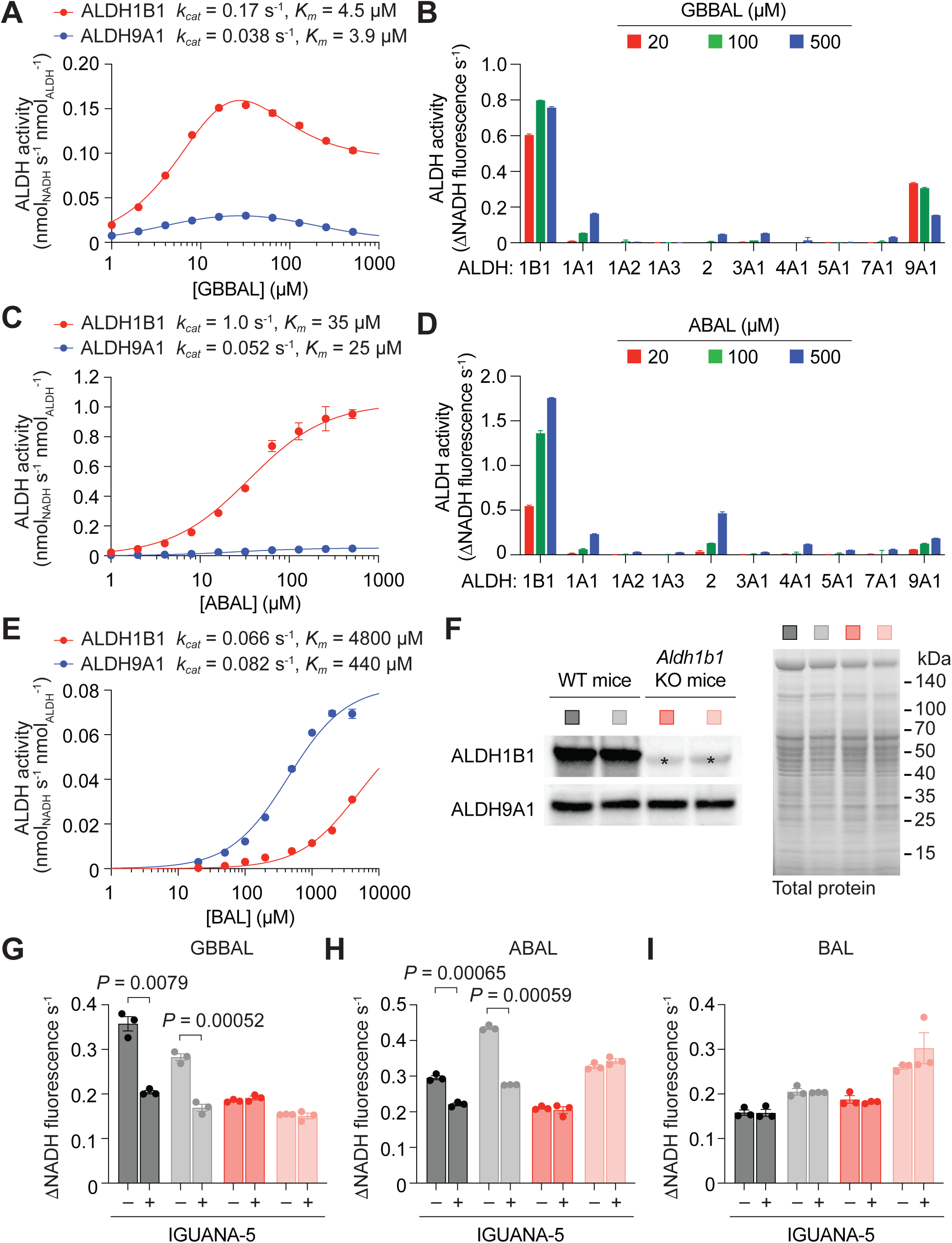
GBBAL and ABAL are preferred substrates of ALDH1B1. (A) Enzymatic activities of recombinant ALDH1B1 and ALDH9A1 using GBBAL as the substrate. (B) Substrate specificity profiling of selected human ALDH isoforms against GBBAL. (C) Enzymatic activities of recombinant ALDH1B1 and ALDH9A1 using ABAL as the substrate. (D) Substrate specificity profiling of selected human ALDH isoforms against ABAL. (E) Enzymatic activities of recombinant ALDH1B1 and ALDH9A1 using BAL as the substrate. ALDH monomer concentrations used in (A), (C), and (E) were 100 nM, and ALDH monomer concentrations used in (B) and (D) were 50 nM. (F) Western blotting analysis of mitochondrial extracts from the livers of wild-type and *Aldh1b1*^−/−^ mice, using ALDH1B1 or ALDH9A1 antibodies. A cross-reactive protein in the anti-ALDH1B1 blot is labeled with an asterisk, and total protein staining for each sample is also shown. (G-I) Enzyme kinetics assays of the wild-type and *Aldh1b1*^−/−^ mitochondrial extracts using 0.1 mM GBBAL (G), 0.2 mM ABAL (H), or 1.0 mM BAL (I) as the substrate and treated with either 2 µM IGUANA-5 or DMSO vehicle alone. Bars for each assay result are colored to match the corresponding mitochondrial extracts shown in (F). All graphed data are the average of three biological replicates ± s.e.m.

To corroborate the results of these recombinant enzyme-based assays, we assessed whether GBBAL and ABAL can be readily metabolized by endogenous ALDH1B1 derived from mammalian cells. We extracted native mitochondrial proteins from the livers of wild-type and *Aldh1b1^−/−^* mice^46^, confirming the absence of ALDH1B1 protein in the latter samples by western blotting (Fig. 3F). Mitochondrial extracts from wild-type mice exhibited GBBAL- and ABAL-oxidizing activities that could be inhibited by IGUANA-5; however, the corresponding extracts from *Aldh1b1^−/−^* mice had only background levels of GBBAL and ABAL oxidation that were IGUANA-5-insensitive (Fig. 3G-H). Consistent with our assay results with purified ALDH1B1 protein, the mitochondrial extracts from wild-type and *Aldh1b1^−/−^* mice oxidized BAL with similar efficacy, and neither activity was inhibited by IGUANA-5 (Fig. 3I).

These observations demonstrate the ability of murine mitochondrial ALDH1B1 to oxidize these aldehyde substrates and independently confirm the specificity of IGUANA-5 for this ALDH family member. We speculate that the IGUANA-5-insensitive GBBAL- and ABAL-oxidizing activities in the mitochondrial extracts could be due to the presence of ALDH9A1 (Fig. 3F), which could reflect cytosolic contamination of these preparations. Alternatively, there may be a mitochondrial form of ALDH9A1 in liver cells, as described in the following section.

### Mitochondrial and cytosolic pools of GBB and GABA are functionally distinct

We next investigated the functional significance of ALDH1B1-dependent GBB and GABA production. GBB is the immediate biosynthetic precursor of carnitine^47,48^, which is conjugated to fatty acids by carnitine palmitoyltransferase 1 (CPT1) to facilitate their transport into mitochondria for oxidative catabolism (Fig. 4A). GBB is produced by the hydroxylation of trimethyllysine, cleavage of the resulting product to form GBBAL and glycine, and GBBAL oxidation. While 3-hydroxytrimethyllysine is generated by the mitochondrial trimethyllysine dioxygenase TMLD^49^, the oxidation of GBBAL to GBB has been attributed to cytosolic ALDH9A1^44^, followed by γ-butyrobetaine dioxygenase 1 (BBOX1)-catalyzed conversion of GBB into carnitine^50^. ALDH9A1 also plays an important role in GABA biosynthesis (Fig. 4B). Although GABA is primarily generated through glutamic acid decarboxylation^51^, the metabolite can be produced through the ALDH9A1-dependent oxidation of ABAL^52^.

**Figure 4.**
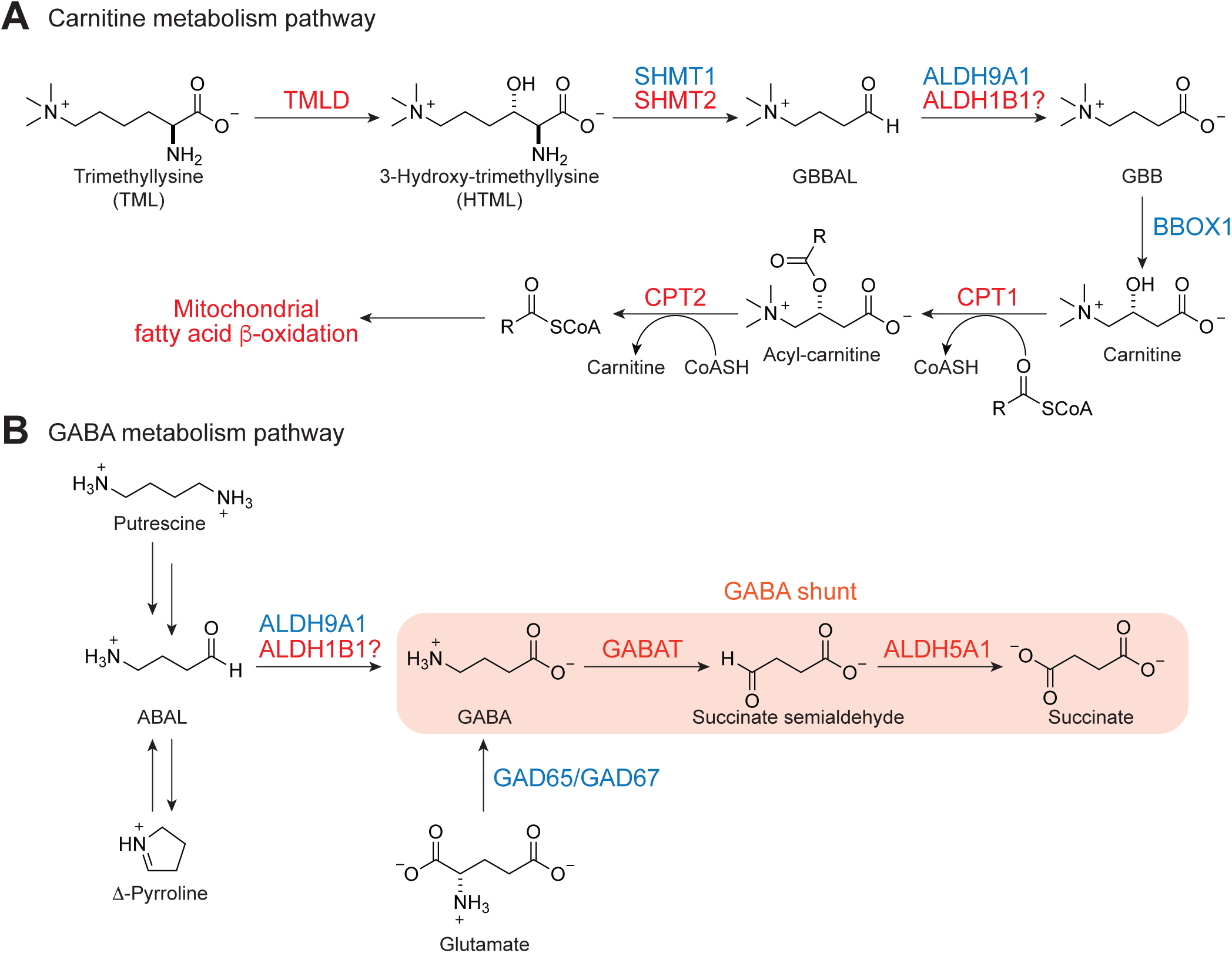
Carnitine and GABA metabolism. (A-B) Enzymatic steps in carnitine (A) and GABA (B) metabolism. Cytosolic and mitochondrial enzymes are shown in blue and red, respectively. Enzyme abbreviations: TMLD, trimethyllysine dioxygenase; SHMT1 and SHMT2, serine hydroxymethyltransferases 1 and 2; BBOX1, γ-butyrobetaine hydroxylase 1; CPT1 and CPT2, carnitine palmitoyltranferases 1 and 2; GABAT, GABA transaminase; GAD65 and GAD67, glutamate decarboxylases 65 and 67.

The results from our metabolomic studies suggest that ALDH1B1 could serve as a mitochondrial counterpart to ALDH9A1. Indeed, the retro-aldol cleavage of 3-hydroxytrimethyllysine into GBBAL and glycine can be mediated by both cytoplasmic and mitochondrial serine hydroxymethyl transferases (SHMT1 and SHMT2, respectively)^53^, suggesting that the GBB could have distinct subcellular activities. GABA is also known to have compartment-specific functions. The metabolite can activate G protein-coupled receptors via presynaptic vesicles^54,55^ or be converted into succinate by the mitochondrial enzymes GABA transaminase (GABAT) and ALDH5A1^56,57^, a pathway commonly referred to as the “GABA shunt.”

We therefore examined whether culturing *ALDH1B1^−/−^* SW480 cells with either metabolite can restore spheroid growth. Neither GBB nor GABA, alone or in combination, could compensate for loss of ALDH1B1 activity (Fig. 5A-C), suggesting that the exogenously applied metabolites cannot substitute for those produced in mitochondria. In fact, exogenous GBB selectively suppressed the viability of ALDH1B1-deficient spheroids (Fig. 5A), suggesting that the balance of mitochondrial and cytosolic GBB influences 3D CRC growth.

**Figure 5.**
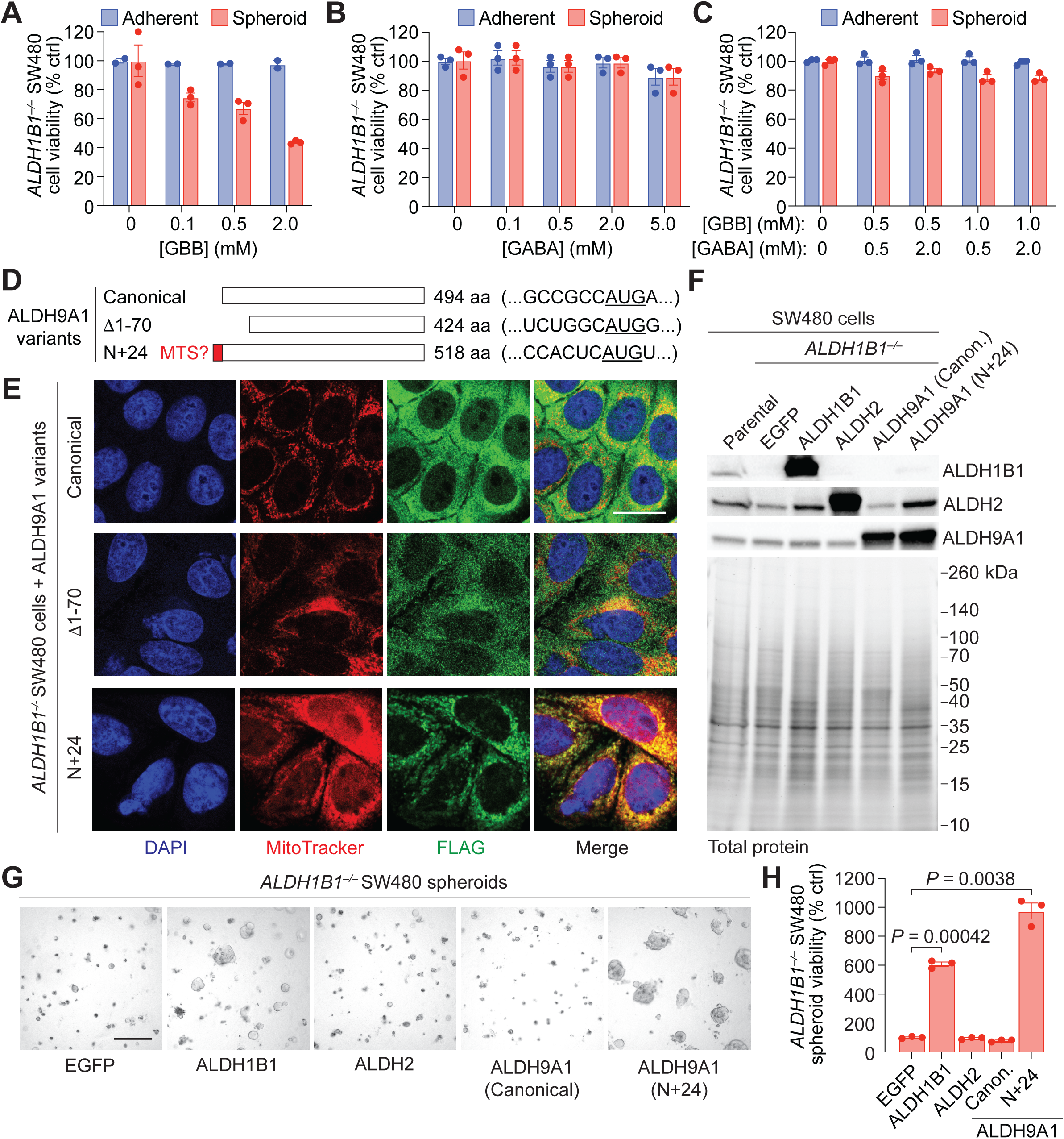
Mitochondrial GBB and/or GABA are required for SW480 spheroid growth. (A-C) Viability of adherent and spheroid cultures of *ALDH1B1*^−/−^ SW480 cells treated with varying concentrations of GBB (A), GABA (B) or both metabolites (C). Data are the average of at least two biological replicates ± s.e.m. and normalized to the vehicle control for each culture condition. (D) The canonical ALDH9A1 isoform and predicted variants. The endogenous Kozak sequence for each isoform is also shown with the start codon underlined. (E) Immunofluorescence images of *ALDH1B1^−/−^* SW480 cells transduced with lentivirus encoding the indicated ALDH9A1 variants with a C-terminal FLAG tag. Mitochondria were labeled with MitoTracker, and nuclei were stained with DAPI. (F-H) Western blot analysis (F), brightfield imaging (G) and viability assays (H) of *ALDH1B1*^−/−^ SW480 cells expressing the indicated FLAG-tagged ALDH isoforms. Data in (H) are the average of three biological replicates ± s.e.m. and normalized to the EGFP control. Scale bars: 20 µm in (E) and 200 µm in (G).

To enable mitochondrial GBB and GABA production in ALDH1B1-deficient cells, we artificially expressed ALDH9A1 in this organelle. Although the canonical form of this enzyme is cytosolic, the UniProt database predicts two additional human isoforms: a variant missing the first 70 amino acids of the canonical form (Δ1-70; produced by alternative splicing) and another with a potential mitochondrial targeting sequence (MTS) (N+24; produced by alternative translation initiation) (Fig. 5D). The canonical and putative Δ1-70 isoform have strong Kozak motifs, whereas the predicted N+24 variant initiates from considerably less favorable sequence^58^. We expressed all three ALDH9A1 variants as C-terminally FLAG-tagged constructs in *ALDH1B1^−/−^* SW480 cells using a CMV promoter and an optimal Kozak sequence. The canonical and Δ1-70 forms were both cytosolic, with the latter construct exhibiting a more punctate distribution. In comparison, the N+24 variant localized to mitochondria (Fig. 5E). These results corroborate a recent report that alternative start codon selection can regulate the subcellular localization of ALDH9A1^59^. We then evaluated the ability of the canonical and N+24 forms of ALDH9A1 to rescue the growth of *ALDH1B1^−/−^*SW480 spheroids. Exogenous ALDH1B1 and ALDH2, another mitochondrial ALDH isoform and the closest structural homolog of ALDH1B1, were used as comparison controls (Fig. 5F and Supplementary Fig. 5). Expressing either ALDH1B1 or the N+24 isoform of ALDH9A1 restored *ALDH1B1^−/−^*spheroid viability, but the canonical ALDH9A1 isoform and ALDH2 did not (Fig. 5G-H). Taken together, these observations suggest that mitochondrial GBB and/or GABA are necessary and sufficient for SW480 spheroid viability.

### ALDH1B1 promotes FAO in CRC spheroids

A role for mitochondrial GBB and carnitine metabolism in CRC would align with recent studies that have associated FAO with this malignancy. Carnitine primarily serves as a fatty acid shuttle for mitochondrial uptake and β-oxidation (Fig. 6A), and FAO activity is frequently elevated in aggressive or metastatic CRC tumors^60–62^. Accordingly, many CRCs exhibit increased *de novo* fatty acid synthesis expression^63,64^, and CPT1 enzymes are upregulated in subsets of CRC cells associated with stem-like and invasive phenotypes^60,61^. FAO has also emerged as an important regulator of stem cell maintenance in the intestine, brain, muscle, and hematopoietic system^65–69^. We therefore focused on our studies on the relationship between ALDH1B1 and FAO.

**Figure 6.**
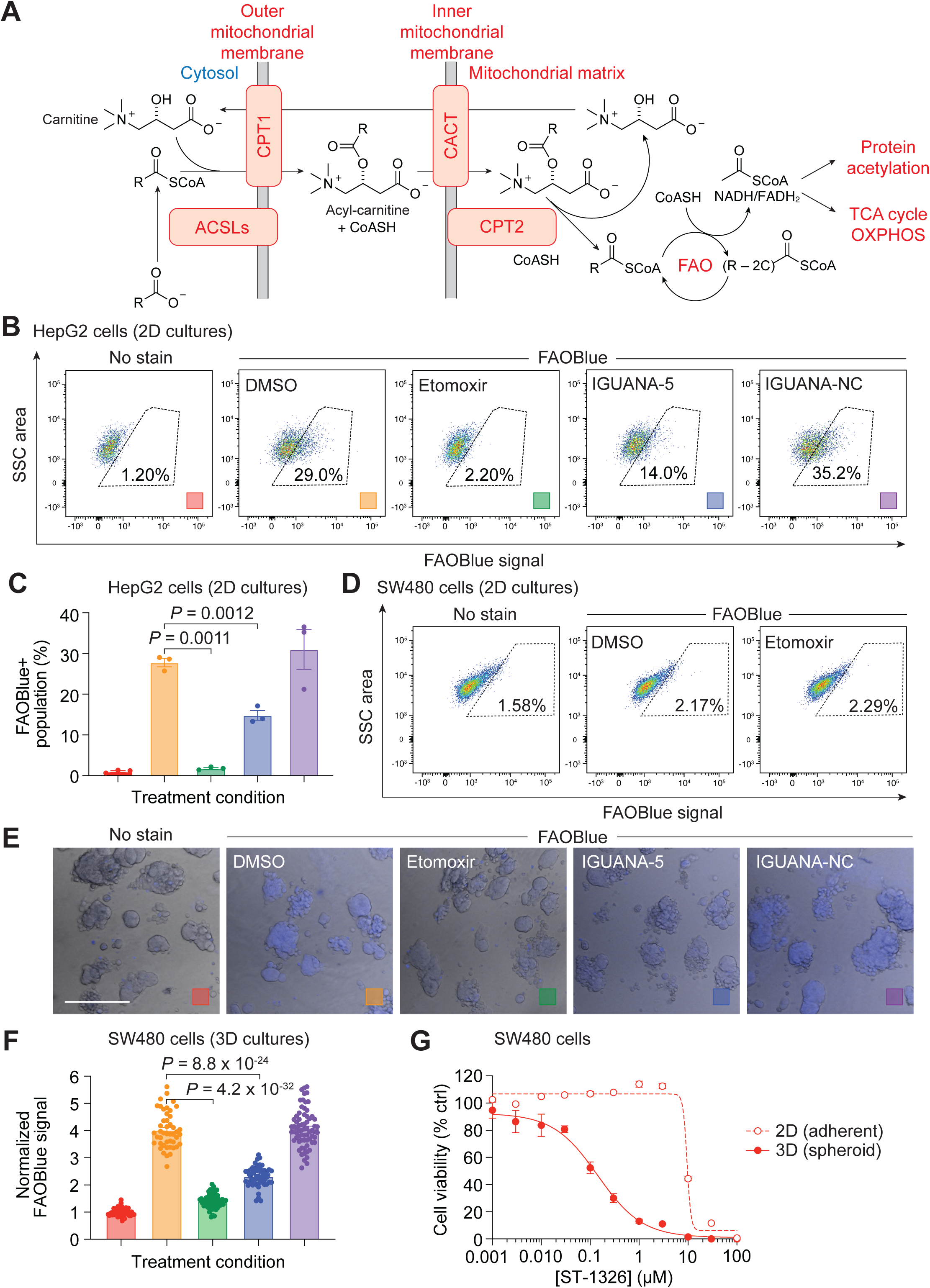
ALDH1B1 promotes FAO in SW480 spheroid cells. (A) Enzymatic and transport steps in mitochondrial fatty acid oxidation. (B) Representative FACS plots for 2D cultures of HepG2 cells treated with DMSO, 10 µM etomoxir, 2 µM IGUANA-5, or 2 µM IGUANA-NC and then and probed with 10 µM FAOBlue dye. 10,000 cells were analyzed for each condition, and FAOBlue-positive populations were gated as shown with the dashed boxes. (C) Quantification of the FACS analyses, with bars colored according to the treatment conditions defined in (B). Data the average of three biological replicates ± s.e.m. (D) Representative FACS plots for 2D cultures of SW480 cells treated with the indicated compounds and FAOBlue as described in (B). (E) Representative images of SW480 spheroids treated with the indicated compounds and FAOBlue. Merged brightfield and fluorescence micrographs are shown. Scale bar: 500 µm. (F) Quantification of the SW480 spheroid micrographs, with bars colored according to the treatment conditions defined in (E). Each data point represents an individual spheroid. (G) Dose-dependent effects of the CPT1 inhibitor ST1326 on 2D and 3D SW480 cell cultures. Data are the average of three biological replicates ± s.e.m.

To determine whether ALDH1B1 influences FAO activity, we used FAOBlue, a fatty acid-mimetic probe that releases a coumarin fluorophore in the final round of β-oxidation^70^. We first examined the effect of IGUANA-5 on FAO in HepG2 hepatoblastoma cells, which express ALDH1B1 and previously have been reported to exhibit detectable FAO activity^70^. For these studies we also prepared a negative control compound, IGUANA-NC, which is inactive in biochemical and cell-based assays of ALDH1B1 activity (Supplementary Fig. 6). We cultured HepG2 cells as 2D monolayers in the presence of IGUANA-5, IGUANA-NC, or DMSO vehicle alone for 48 h. The cells were then treated with the FAOBlue probe for 48 h and then analyzed by flow cytometry. HepG2 cells cultured with the covalent CPT1 inhibitor etomoxir for 4 h prior to FAOBlue treatment were used to establish the background coumarin fluorescence in HepG2 cells lacking FAO activity (Fig. 6B-C). We observed that ALDH1B1 inhibition decreased FAO activity in the liver cancer cells by about two-fold, while the control compound IGUANA-NC had no effect.

We next examined whether ALDH1B1 also promotes FAO in SW480 CRC cells. When maintained as 2D monolayers, the SW480 cells exhibited minimal FAOBlue fluorescence, even with elevated concentrations of the FAO probe (Fig. 6D and Supplementary Fig. 7A). However, we observed robust etomoxir-sensitive FAOBlue fluorescence in SW480 spheroids, which we could quantify through confocal microscopy (Fig. 6E-F). As in our HepG2 experiments, the FAO activity in these 3D CRC cultures was suppressed two-fold by IGUANA-5 but unaffected by IGUANA-NC. We also assessed the dependency of SW480 cells on FAO by treating 2D and 3D cultures of this CRC line with the reversible CPT1 inhibitor ST1326 (Fig. 6G). The FAO-blocking compound preferentially suppressed the growth of SW480 spheroids, recapitulating the effects of IGUANA-5 (Supplementary Fig. 6C).

Collectively, these findings establish FAO as a metabolic dependency of SW480 spheroids and a role for ALDH1B1 in achieving the level of FAO activity required for CRC growth. In contrast, adherent SW480 cultures are markedly less reliant on FAO and ALDH1B1 for their viability. To explore the potential mechanisms underlying the divergent FAO dependencies of 2D and 3D SW480 cultures, we used RNA sequencing to compare the transcriptomes associated with the two growth conditions. Among the genes encoding enzymes that contribute to mitochondrial FAO, we found that *ACSL5*, a member of the acyl-CoA synthetase long-chain family, is markedly upregulated in SW480 spheroids (Supplementary Fig. 7B and Supplementary File 2). Thus, the elevated FAO activity in these 3D cultures could stem, at least in part, from higher levels of the fatty acid-CoA substrates for CPT1.

### GBB functions as a substrate for the mitochondrial transporter CACT

We concluded our studies by investigating how ALDH1B1 might drive FAO in CRC, focusing on the role of this enzyme in GBB production. Since the primary role of GBB in mammals has been ascribed to its conversion to carnitine by the cytosolic enzyme BBOX1, ALDH1B1 might enhance FAO by increasing acylcarnitine levels. However, the specific requirement of mitochondrial GBB in SW480 spheroids suggests that ALDH1B1-generated GBB modulates FAO through a different mechanism. We hypothesized mitochondrial GBB could drive CACT-mediated acylcarnitine uptake, which has two modes of action: uniport of acylcarnitine into mitochondria and carnitine-acylcarnitine antiport. The antiport mechanism has been estimated to be at least 10-fold faster than uniport^71,72^, and the mitochondrial carnitine used in this process is derived from the carnitine palmitoyltransferase 2 (CPT2)-mediated conversion of imported acylcarnitines into their corresponding acyl-CoAs. This functional framework creates an apparent circular dependency: efficient acylcarnitine transport into mitochondria requires mitochondrial carnitine, whereas mitochondrial carnitine is itself derived from imported acylcarnitines. However, if GBB can serve as a carnitine surrogate due to their structural similarity, ALDH1B1 could provide an independent mechanism for maximizing FAO activity to maintain stemness and promote cancer growth.

To test this hypothesis, we utilized an *in vitro* proteoliposome-based assay for CACT activity (Fig. 7A). In this method, purified recombinant human CACT is reconstituted into L-α-phosphatidylcholine liposomes preloaded with defined internal substrates. CACT uniport and antiport activities can then be measured by quantifying the uptake of external [^3^H]-carnitine. CACT-harboring proteoliposomes with luminal carnitine efficiently transported external [^3^H]-carnitine as previously reported (Fig. 7B-C). Proteoliposomes loaded with GBB also supported robust [³H]-carnitine uptake at levels comparable to that mediated by luminal carnitine. In contrast, betaine showed minimal activity that was indistinguishable from buffer-only controls, consistent with previous studies demonstrating that this metabolite is not a CACT substrate.^73–75^ These data establish GBB as a bona fide substrate for CACT that can functionally substitute for carnitine as an antiport substrate. Together with our ALDH9A1 rescue experiments, these findings support a model in which ALDH1B1-generated GBB within the mitochondrial matrix directly facilitates acylcarnitine import via CACT, thereby driving FAO to the levels necessary to sustain stem-like CRC cells.

**Figure 7.**
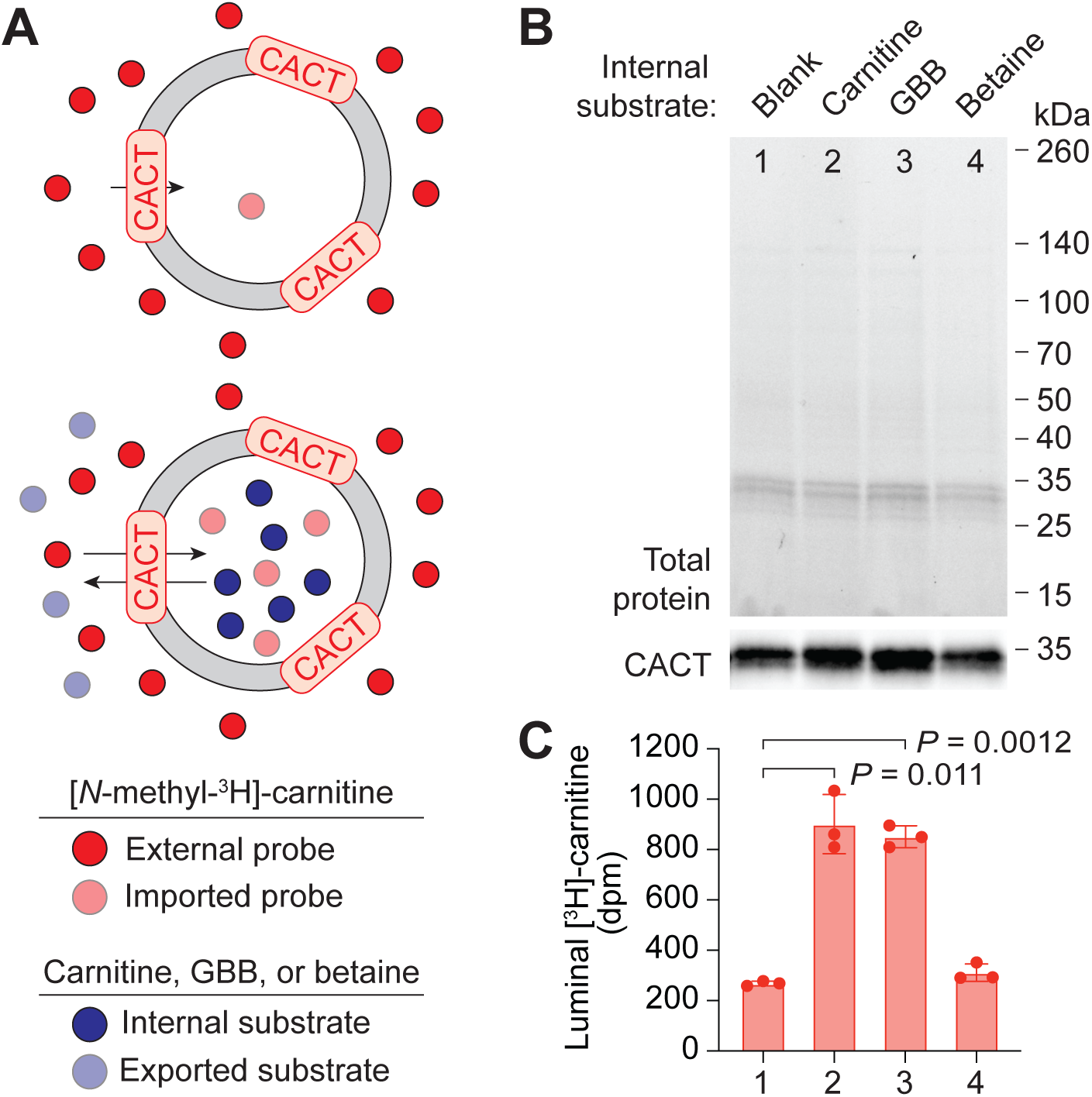
GBB functions as a CACT substrate. (A) Schematic of the proteoliposome-based assay for CACT-mediated antiport. (B) Total protein staining and anti-CACT western blot analyses of proteoliposomes reconstituted with CACT and preloaded with buffer (blank) or the indicated internal substrates. (C) [^3^H]-carnitine uptake by the substrate-preloaded CACT proteoliposomes, Bar numbers correspond to the internal substrates defined in (B), and data are the average of three biological replicates ± s.e.m.

## DISCUSSION

High ALDH1 activity is a well-established marker of normal stem and progenitor cells, and it has emerged as a characteristic feature of stem-like cells in multiple cancers. As individual ALDH1 isoforms gain prominence as potential therapeutic targets, elucidating how these enzymes support stemness and tumor growth has become an important focus of ongoing research. Early studies explored the association between high ALDH1 expression and chemoresistance, leading to the discovery that ALDH1A1 can detoxify cyclophosphamide and other oxazaphosphorine-based alkylating chemotherapies by oxidizing the pro-drug intermediate aldophosphamide^76–78^. More recent investigations have focused on the intrinsic roles of ALDH1 enzymes in CSC maintenance and tumor progression, including their endogenous substrates and contributions to cellular metabolism and signaling^79–81^. For example, ALDH1A3 has been shown to coordinate nuclear acetaldehyde/acetyl-CoA metabolism with histone H3 acetylation, leading to the expression of neural crest stem cell and glucose metabolism genes in melanoma^14^.

Here we establish a molecular and cellular framework for understanding ALDH1B1 function. While a recent study proposed that this mitochondrial ALDH1 isoform protects cells from ferroptosis^32^, our findings indicate that lipid aldehyde detoxification is unlikely to be the primary means by which ALDH1B1 promotes CRC. ALDH1B1 inhibition does not increase the sensitivity of CRC cells to ferroptosis inducers, and ferroptosis inhibitors do not restore the viability of *ALDH1B1^−/−^* CRC spheroids. Moreover, these ALDH1B1-deficient 3D cultures cannot be rescued by overexpressing ALDH2, an ALDH isoform known to detoxify lipid peroxidation products and suppress ferroptosis.^82,83^ It has also been reported that ALDH1B1 upregulates Wnt, Notch, and PI3K/Akt signaling in CRC cells; however, the mechanistic relationship between ALDH1B1 and these oncogenic pathways remains unknown and is probably indirect. Our work provides a more direct assessment of ALDH1B1 function in CRC, leveraging mass spectrometry-based metabolomics and the temporal control afforded by ALDH1B1-specific small molecules. We establish GBBAL and ABAL as preferred substrates of ALDH1B1, uncovering a mechanism for the mitochondrial production of GBB and GABA, respectively.

The preference of ALDH1B1 for GBBAL and ABAL substrates is consistent with structural features of its active site. ALDH1B1 originated from a retrotransposition of ALDH2^84^, and the two paralogs share 72% sequence identity. Phylogenetic analyses have identified *ALDH1B1* genes in several mammalian species and in frogs^84^. However, no ALDH1B1 homologs have been found in the genomes of birds, reptiles, or fish, indicating that this mitochondrial enzyme is a relatively recent addition to the ALDH family. ALDH1B1 retains the ability of ALDH2 to oxidize acetaldehyde^85,86^, and its reactivity toward GBBAL and ABAL coincides with the presence of several acidic residues in the active site (Glu116, Asp121, Asp123, and Glu124; residue numbering is based on the mature mitochondrial protein) (Supplementary Fig. 8). This cluster of negatively charged amino acids could complement the cationic quaternary amine in GBBAL and protonated amine in ABAL, and it is unique within the ALDH family. ALDH9A1 has three of the four Asp/Glu residues found in ALDH1B1, and the other ALDH isoforms have only one or two. In addition, all four acidic residues are conserved in plant ALDH10 enzymes, which are aminoaldehyde dehydrogenases with preferential activity toward GBBAL and ABAL. Consistent with our findings, *Arabidopsis* lines lacking *aldh10a8* and *aldh10a9* also have impaired GBB and GABA production^87,88^. Thus, the evolution of ALDH1B1 into a GBB- and GABA-producing enzyme appears to be an example of functional convergence.

The capacity of ALDH1B1 to produce GBB and GABA also highlights subcellular compartmentalization as a mechanism for regulating metabolite function. For this study, we have focused on the ALDH1B1-GBB axis, revealing compartment-specific functions of cytosolic and mitochondrial GBB (Fig. 8). Cytosolic GBB is a precursor for carnitine biosynthesis, driving FAO through the CPT1-CACT-CPT2 fatty acid transport cycle. Our findings support a model in which mitochondrial GBB promotes FAO activity through at least two mechanisms. First, GBB within the mitochondrial inner matrix replaces carnitine as a CACT antiport substrate, circumventing the reciprocal coupling of acylcarnitine uptake and carnitine production in this organelle. Second, CACT-transported GBB likely crosses the mitochondrial outer membrane and augments the cytosolic GBB pool that is converted into carnitine, which could account for the lower carnitine and acylcarnitine levels we observed in ALDH1B1 inhibitor-treated SW480 cells. These compartment-specific GBB functions are likely reinforced by the presence of both cytosolic and mitochondrial SHMT isoforms, which are believed to catalyze the retro-aldol cleavage of HTML into GBB and glycine. Our studies further indicate that reprogramming GBB biosynthesis from the cytosol to mitochondria has a sizable effect on FAO flux, as endogenous ALDH1B1 expression approximately doubles the FAO activity in hepatoblastoma and CRC cells.

**Figure 8.**
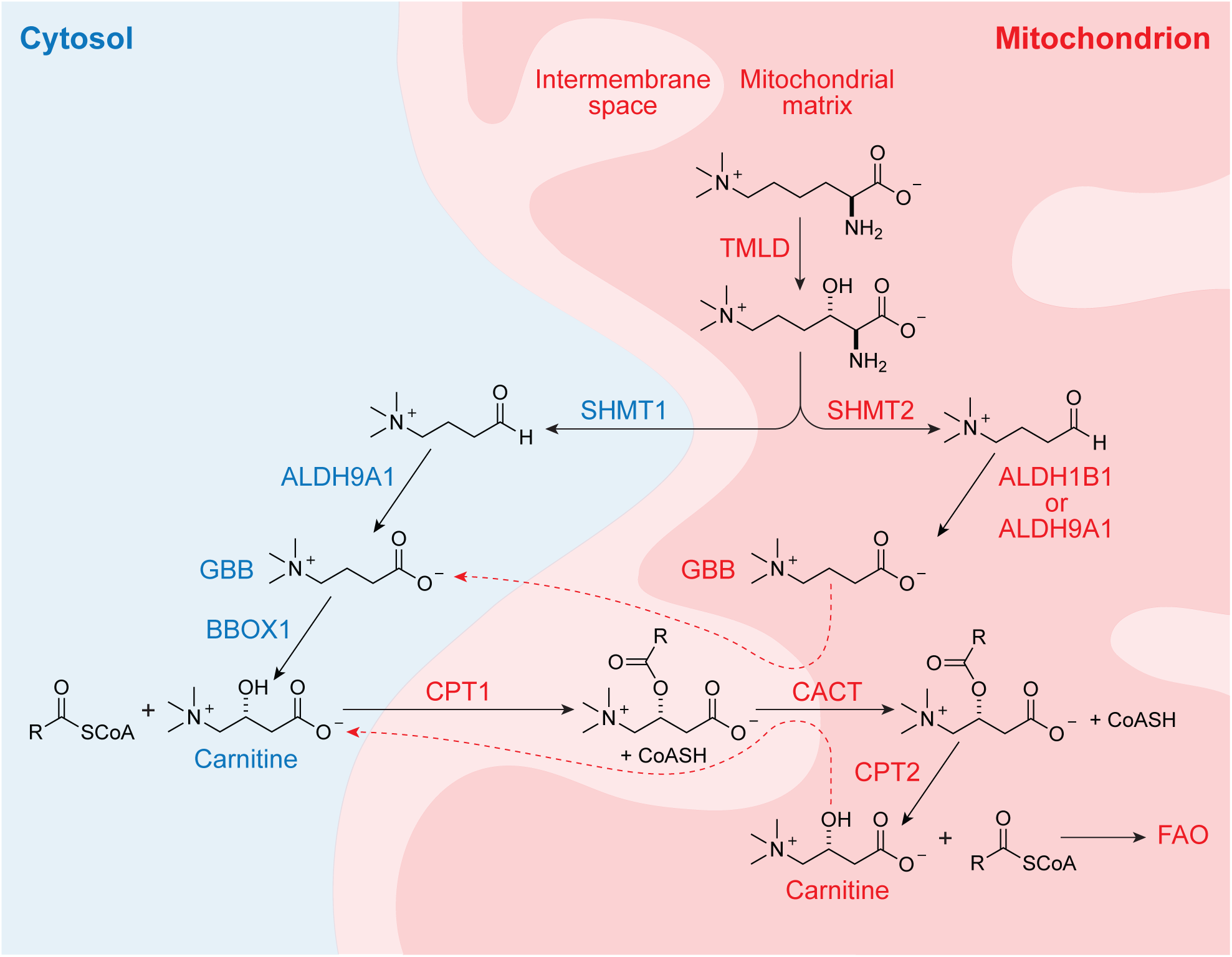
A model for compartment-specific roles of GBB in FAO regulation. Parallel pathways for GBB biosynthesis exist in the cytosol and mitochondrial inner matrix, catalyzed respectively by SHMT1/ALDH9A1 or SHMT2/ALDH1B1 axes. It is also possible that a mitochondrial variant of ALDH9A1 promotes GBB production in this subcellular compartment in certain cellular contexts. Cytosolic GBB supports carnitine production, whereas mitochondrial GBB enhances acylcarnitine transport into via CACT, a process that is normally mediated by mitochondrial carnitine. Like carnitine, GBB that enters the intermembrane space via CACT could also transit the outer mitochondrial membrane, contributing to the cytosolic GBB pool that is oxidized by BBOX1. Mitochondrial GBB therefore both uncouples acylcarnitine uptake and hydrolysis and supports carnitine biosynthesis, enabling cancer cells to sustain elevated FAO activities that support stemness and tumor growth.

Previous investigations have established FAO as a metabolic requirement of intestinal stem cells,^67,68,89,90^ and fatty acid catabolism promotes CRC cell proliferation, metastasis, and stemness.^60,62,68^ Earlier work by our lab also showed that ALDH1B1 sustains the expression of genes associated with CRC stem cells^23^. Thus, our new findings establish ALDH1B1 as a rheostat of FAO activity that affords mammals and amphibians a yet-to-be-determined fitness advantage and has been co-opted by CRC to sustain stem-like tumor cells. ALDH1B1 may act through analogous mechanisms in PDAC, osteosarcoma, nasopharyngeal carcinoma, and other cancers linked to this enzyme^91,92^.

Other examples of compartment-specific metabolic biosynthesis and function include the mitochondrial and nucleocytosolic production of acetyl-CoA by the synthases ACSS1 and ACSS2, respectively, which diverged from a common ancestral enzyme. Acetyl-CoA generated by ACSS1 enters the TCA cycle^93^, whereas the same metabolite derived from ACSS2 supports lipid synthesis^94^. Compartmentalized NAD^+^ synthesis by nuclear and cytosolic nicotinamide mononucleotide adenylyl transferases (NMNATs) similarly enables reciprocal control of NAD^+^-dependent gene regulation and cellular metabolism^95^.. A single gene can also enable the reprogramming of metabolite production to specific subcellular compartments through alternative splicing, transcription initiation, or translation start-site selection. For example, the long-chain acyl-CoA synthetase ACSL6 can be expressed as four catalytically active isoforms that vary in active site and N-terminal sequences, two of which preferentially accept docosahexaenoic acid as substrates but exhibit distinct subcellular distributions^96^. Our results raise the possibility that ALDH9A1 variants could be analogously deployed to spatially regulate GBB biosynthesis.

The ability of ALDH1B1 to shift GBB and GABA production from the cytosol to mitochondria represents a third paradigm of spatial metabolic rewiring. In this case, subcellular control has been achieved through the evolution of a mitochondrial ALDH enzyme to biochemically emulate a distantly related paralog with predominantly cytosolic localization. Despite their convergent substrate preferences, ALDH1B1 and ALDH9A1 share only 39% sequence identity, and the two enzymes afford orthogonal control of carnitine metabolism and FAO. Indeed, the minimal effects of ALDH1B1 deficiency on mammalian development and physiology might be explained by upregulation of mitochondrial ALDH9A1. Conversely, the pronounced effects of genetically or pharmacologically abrogating ALDH1B1 activity on CRC and PDAC suggest that these cancers have more limited plasticity with respect to ALDH9A1 function. Given the therapeutic potential of ALDH1B1 inhibitors, the cellular pathways that regulate *ALDH9A1* translation initiation and their propensity to be co-opted as drug resistance mechanisms are important areas for future investigation. Inhibitors that selectively target mitochondrial ALDH9A1 might also have therapeutic potential.

Finally, our findings highlight areas for future mechanistic studies. For example, ALDH1B1-driven FAO should increase mitochondrial levels of NADH, FADH_2_, and acetyl-CoA, and how these metabolites promote CRC remain to be determined. In principle, the higher FAO activity could lead to increases in TCA cycle flux, protein acetylation, or both. The relative contribution of ALDH1B1-mediated GABA production to CRC is another open question. The “GABA shunt” converts GABA into succinate, a known oncometabolite. Alternatively, GABA has been reported to act through the GABA_B_ receptor and β-catenin to activate the Wnt pathway^97–99^, a major driver of CRC^100^. An ability of ALDH1B1 to potentiate both FAO and Wnt pathway activity could underlie its role in CRC, PDAC, and other human malignancies. Finally, the functional consequences of ALDH1B1 deficiency could extend beyond the loss of mitochondrial GBB and GABA. In addition to the possibility of other direct ALDH1B1 substrates and products, mitochondrial accumulation of GBBAL or ABAL could preferentially disrupt enzymes that act on structurally related metabolites. Such a mechanism could account for the reduced betaine levels in ALDH1B1-treated SW480 cells. Small-molecule inhibitors of ALDH1B1 and other enzymes will be valuable tools for delineating these metabolic and signaling pathways.

## METHODS

### Chemicals

Acetaldehyde, all-trans retinal, DEAB, NAD^+^, BAL (betaine aldehyde chloride), GBB (**γ−**butyrobetaine hydrochloride), GABA (**γ−**aminobutyric acid), and NEM (N-Ethylmaleimide) were purchased from Sigma Aldrich. 4-HNE, RSL3, Ferrostatin-1 and Liproxstatin-1 was purchased from MedChemExpress. Carnitine (L(-)-Carnitine hydrochloride) and DTE (dithioerythritol) were purchased from Thermo Scientific. [^3^H]-carnitine (L-Carnitine [N-methyl-3H] hydrochloride) was purchased from CliniSciences. ABAL was prepared from 4-aminobutyraldehyde diethyl acetal (Combi-Blocks) according to the method described by Hanson and coworkers^101^.

GBBAL was synthesized by the following modification of a previous protocol^44^, yielding a solid iodide salt that could be stably stored under an inert atmosphere at room temperature. A flask was charged with 4,4-diethoxy-*N,N*-dimethyl-1-butanamine (0.40 mL, 1.8 mmol, 1.0 eq.), acetonitrile (3 mL), and 1 M HCl in water (2 mL). The reaction was stirred at room temperature for 3 h and then transferred to a separatory funnel and diluted with water (5 mL). The solution was washed with dichloromethane (2 x 5 mL), and the aqueous layer was basified with Na_3_PO_4_ (0.5 g, 3 mmol) and then extracted with dichloromethane (2 x 5 mL). The organic layers were combined, dried over Na_2_SO_4_, filtered, and concentrated under reduced pressure (80 mbar, 22 °C, 10 min). The crude residue was diluted with chloroform (5 mL) and split evenly between two 20-mL vials. Each vial was then charged with iodomethane (0.08 mL, 1.3 mmol, 1.5 eq.), sealed under N_2_ and gently agitated at room temperature for 1 h. The reaction was next concentrated under reduced pressure (<10 mbar, 22 °C, 5 min), and the resulting solid was dried in a vacuum desiccator for 16 h to afford the desired product as a white, crystalline solid (0.080 g, 0.31 mmol, 17% yield). ^1^H NMR (500 MHz, DMSO-*d_6_*) δ 9.68 (t, *J* = 1.0 Hz, 1H), 3.28 – 3.23 (m, 2H), 3.06 (s, 9H), 2.57 (td, *J* = 7.0, 1.0 Hz, 2H), 1.92 (p, *J* = 7.8 Hz, 2H). ^13^C NMR (126 MHz, DMSO-*d_6_*): δ 201.93, 64.45 (t, *J* = 3.1 Hz), 52.17 (t, *J* = 4.0 Hz), 15.18. HRMS (H-ESI) *m/z*: [M]^+^ calcd for C_7_H_16_NO 130.1226; found 130.1225.

IGUANA-5 was synthesized as previously described^33^. IGUANA-NC was synthesized in six steps from 3-phenylcylobutan-1-one according to the following procedure:

#### N-(3-phenylcyclobutylidene)hydroxylamine

To a solution of hydroxylamine hydrochloride (3.14 g, 45.2 mmol, 2.2 eq.) in methanol (15 mL) and water (1.65 mL) was added K_2_CO_3_ (6.24 g, 45.2 mmol, 2.2 eq.). After 10 min, 3-phenylcyclobutan-1-one (3.0 g, 20.5 mmol, 1.0 eq.) was added, and the resulting reaction mixture was heated to 50 °C for 14 h, The reaction mixture was next diluted with ethyl acetate (100 mL) and washed sequentially with water (50 mL) and brine (40 mL). The organic layer was dried over Na_2_SO_4_, filtered, and concentrated under reduced pressure to obtain the desired product as a brown liquid (3.0 g, 18.6 mmol, 90% yield). ^1^H NMR (400 MHz, DMSO-*d*_6_): δ 10.32 (s, 1H), 7.37–7.25 (m, 4H), 7.23-7.17 (m, 1H), 3.63–3.46 (m, 1H), 3.38–3.17 (m, 2H), 2.92-2.76 (m, 2H).

#### 3-phenylcyclobutan-1-amine

To a solution of *N*-(3-phenylcyclobutylidene)hydroxylamine (4.0 g, 24.8 mmol, 1.0 eq.) in methanol (40 mL) in a hydrogenation vessel was added Raney-Ni (11.5 g, 134 mmol, 5.4 eq.), followed by NH_3_ in methanol (40 mL. The resulting reaction mixture was hydrogenated at 2.5 kg/cm^2^ H_2_ pressure for 2 h. The reaction was next filtered through a Celite bed, and the filtrate was concentrated under reduced pressure to obtain the desired product as an oil (3.0 g, 20.4 mmol, 82% yield). ^1^H NMR (400 MHz, CDCl_3_): δ 7.37–7.23 (m, 3H), 7.25–7.14 (m, 2H), 3.76–3.52 (m, 1H), 3.07 (tt, *J* = 10.3, 7.6 Hz, 1H), 2.87–2.66 (m, 2H), 2.57–2.38 (m, 1H), 2.32–2.12 (m, 1H), 1.97–1.71 (m, 2H). LC-MS (ESI) *m/z*: [M+H]^+^ calcd for C_10_H_14_N 148.1; found 148.1.

#### 3-isothiocyanatocyclobutyl)benzene

To a solution of 3-phenylcyclobutan-1-amine (3.1 g, 21.1 mmol, 1.0 eq.) in dichloromethane (31 mL) was added thiophosgene (1.76 mL, 23.2 mmol, 1.1 eq.). After stirring for 2 h at room temperature, the mixture was diluted with dichloromethane (100 mL) and washed with water (50 mL). The aqueous layer was separated and extracted with dichloromethane (50 mL), and the combined organic layers were washed with brine (30 mL), dried over Na_2_SO_4_, filtered, and concentrated under reduced pressure. The crude material was purified by silica-gel column chromatography (60-120 mesh) to obtain the desired product as a pale-yellow liquid (2.9 g, 15.3 mmol, 72% yield). ^1^H NMR (400 MHz, CDCl_3_): δ 7.39-7.31 (m, 2H), 7.29–7.18 (m, 3H), 4.09 (tt, *J* = 9.1, 7.3 Hz, 1H), 3.41–3.10 (m, 1H), 2.95-2.83 (m, 1H), 2.72 (ddd, *J* = 12.2, 9.3, 5.0 Hz, 1H), 2.68–2.56 (m, 1H), 2.40 (dtd, *J* = 11.5, 8.9, 2.8 Hz, 1H).

#### N-(3-phenylcyclobutyl)-1,4,5,6-tetrahydropyrimidin-2-amine

To a solution of 3-isothiocyanatocyclobutyl)benzene (800 mg, 4.23 mmol, 1.0 eq.) in tetrahydrofuran (10 mL) was added propane-1,3-diamine (313 mg, 4.23 mmol, 1.0 eq.). The resulting mixture was heated to 85 °C for 12 h, and then (diacetoxyiodo)benzene (2.72 g, 8.46 mmol, 1.0 eq.) was added to the reaction mixture. After an additional 12 h, the reaction was removed from the heat, and the tetrahydrofuran was evaporated away. The crude material was diluted with ethyl acetate (50 mL), and washed with H_2_O (30 mL), and the organic layer was dried over Na_2_SO_4_, filtered, and concentrated under reduced pressure. The crude material was purified by neutral alumina column chromatography to afford the desired product as a brown solid (450 mg, 46% yield). LC-MS (ESI) *m/z*: [M+H]^+^ calcd for C_14_H_20_N_3_ 230.2; found 230.1.

#### 3-(2-methoxyphenyl)-1-(3-phenylcyclobutyl)-1,5,6,7-tetrahydroimidazo[1,2-a]pyrimidine trifluoroacetic acid salt (IGUANA-NC)

A flask was charged with *N*-(3-phenylcyclobutyl)-1,4,5,6-tetrahydropyrimidin-2-amine (0.500 g, 2.18 mmol, 1.0 eq.) and *N*,*N*-dimethylformamide (5 mL), and cooled to 0 °C. The solution was then treated with 60% NaH in mineral oil (131 mg, 3.27 mmol, 1.0 eq.) and stirred at 0 °C for 30 min, after which 2-bromo-1-(2-methoxyphenyl)ethan-1-one (749 mg, 3.27 mmol, 1.5 eq.) was added. The reaction mixture was warmed to room temperature and after 16 h, it was quenched with saturated aq. NH_4_Cl (20 mL) and extracted with dichloromethane (2 x 60 mL). The combined organic layers were dried over Na_2_SO_4_, filtered, and concentrated under reduced pressure to obtain a brown oil. The residue was dissolved in diphenyl ether (10 mL), treated with added Et_3_N (1.2 mL, 8.73 mmol, 4.0 eq.), and heated to 130 °C for 16 h. The crude was then purified by reverse-phase HPLC to obtain the desired product as an off-white solid (50 mg, 0.14 mmol, 6.3% yield). ^1^H NMR (400 MHz, DMSO-*d_6_*): δ 7.95 (s, 1H), 7.63–7.43 (m, 1H), 7.33 (dq, *J* = 6.7, 2.9, 2.3 Hz, 1H), 7.25–7.17 (m, 2H), 7.17–7.09 (m, 3H), 7.05 (t, *J* = 7.4 Hz, 1H), 7.01–6.90 (m, 2H), 4.47 (ddt, *J* = 10.5, 7.8, 4.3 Hz, 1H), 3.98 (t, *J* = 5.6 Hz, 2H), 3.89–3.71 (m, 3H), 3.43 (s, 2H), 3.07 (t, *J* = 9.1 Hz, 1H), 2.55-2.48 (m, 2H), 2.19–1.95 (m, 4H). ^13^C NMR (126 MHz, DMSO-*d_6_*): δ 157.72, 143.29, 142.83, 131.82, 131.78, 128.25, 126.36, 126.33, 123.99, 120.72, 116.57, 114.56, 111.24, 55.51, 46.56, 42.84, 38.38, 35.98, 32.60, 19.44. HRMS (H-ESI) *m/z*: [M+H]^+^ calcd for C_23_H_26_N_3_O 360.2070; found 360.2066.

### Cell lines

SW480, HT-1080, HepG2 and HEK-293T cells were purchased from ATCC and maintained in RPMI-1640 (SW480) or DMEM media (HT-1080, HepG2 and HEK-293T) containing 10% fetal bovine serum (FBS; Millipore Sigma), 100 U mL^−1^ penicillin and 100 µg mL^-1^ streptomycin. ALDH1B1-overexpressing *ALDH1A3*^−/−^ A375 melanoma cells^33^ were maintained in DMEM containing 10% FBS, 100 U mL^−1^ penicillin, 100 µg mL^−1^ streptomycin and 1.0 µg mL^−1^ puromycin. HT-1080^N^ cells^102^ were maintained in DMEM containing 10% FBS, 100 U mL^−1^ penicillin, 100 µg mL^−1^ streptomycin and 1.5 µg mL^−1^ puromycin.

The *ALDH1B1^−/−^* SW480 clonal cell line^23^ was maintained in RPMI-1640 containing 10% FBS, 100 U mL^−1^ penicillin and 100 µg mL^−1^ streptomycin. To generate derivatives expressing EGFP or exogenous ALDH enzymes, the indicated ALDH1B1, ALDH2, and ALDH9A1 sequences were amplified by PCR using cDNA obtained from the parental SW480 cell line. cDNA encoding C-terminal 3×FLAG epitope tag was introduced into each gene, and the resulting fragments were cloned into the pCDH-CMV-EF1-Puro vector via Gibson Assembly (NEB, E2611S). Individual constructs were validated by whole-plasmid sequencing, and lentiviruses were separately produced by transfecting the individual expression plasmids into HEK-293T cells along with third-generation packaging plasmids (Addgene, 12251, 12253, and 12259). *ALDH1B1^−/−^* SW480 cells were then transduced with the respective lentiviruses, and stable cell lines were generated through puromycin selection (2.0 μg mL^-1^) and maintained in RPMI-1640 containing 10% FBS, 100 U mL^−1^ penicillin, 100 µg mL^−1^ streptomycin and 2.0 µg mL^−1^ puromycin. Lentiviral experiments performed for this study comply with the Stanford University Institutional Biosafety Committee regulations (Stanford IBC #5565).

### Cell viability assays

HT-1080 cells were seeded into 96-well plates at a density of 10,000 cells/well and cultured in media containing RSL with DMSO, IGUANA-5, or DEAB at the indicated concentrations for 48 h. Cell viabilities were then measured using a CellTiter-Glo 3D Cell Viability Assay kit (Promega) and a Veritas microplate luminometer (Turner BioSystems). As an alternative approach for assessing HT-1080^N^ cell viability, the cells were diluted to a density of 25,000 cells/mL and plated in 96-well plates at a density of 5,000 cells in 200 µL culture media per well. The next day, media was removed from each well and replaced with media containing DMSO or IGUANA-5 of the appropriate concentrations for 24 h. The media was then replaced again with media containing RSL3 and DMSO or IGUANA-5, and the cells were imaged 48 h later in a Sartorius Incucyte S3 (Essen Biosciences). All media contained 20 nM SYTOX green (Life Technologies) to quantify dead cells. A lethal fraction score was calculated using the scalable time-lapse analysis of cell death kinetics (STACK) as previously described^102^.

To assess the viability of adherent SW480 cultures, parental SW480 cells, *ALDH1B1^-/-^* SW480 cells and *ALDH1B1^-/-^*SW480 cells overexpressing EGFP or indicated ALDHs were seeded into 96-well plates at 2,000 cells/well. Corresponding SW480 spheroid cultures were established using pre-chilled 96-well plates coated with 25 µL of ice-cold Matrigel per well and incubated at 37 °C for 30 min to allow for matrix polymerization. SW480 cells were suspended in a defined spheroid culture medium composed of RPMI-1640 supplemented with 20 ng mL⁻¹ recombinant human epidermal growth factor (Abcam, ab9697), 20 ng mL⁻¹ recombinant human fibroblast growth factor (BioVision, 4037), 5 µg mL⁻¹ heparin (Sigma), 1× N-2 supplement (Thermo Fisher Scientific), 100 U mL⁻¹ penicillin, and 100 µg mL⁻¹ streptomycin. Cells were then seeded onto the polymerized Matrigel at 2,000 cells/well. Experimental compounds diluted in the same spheroid culture medium were added to the wells, and the cultures were maintained at 37 °C in a humidified 5% CO₂ atmosphere for 4 or 7 days, as specified. Cell viabilities were then measured using the CellTiter-Glo 3D Cell Viability Assay kit Nand Veritas microplate luminometer.

### Metabolomics analysis

*ALDH1B1^-/-^* SW480 cells overexpressing either ALDH1B1 or EGFP were seeded into 6-well plates at a density of 3.0 × 10^5^ cells/well in 3.0 mL of culture medium. Each experimental condition was performed with five biological replicates. After 24 h, the cells were treated with DMSO, 2 µM IGUANA-5 or 2 µM IGUANA-NC for 12 h. The cells were then washed twice on ice with 2 mL of ice-cold 150 mM ammonium acetate (pH 7.3) to remove residual media. Metabolites were extracted by adding 500 µL of chilled 80% (v/v) methanol in water with 10 nM trifluoromethanesulfonate (TFMS) to each well and incubating plates at -80 °C for 15 min. The plates were placed on ice, and the cells were scraped into the methanol solution and transferred to microcentrifuge tubes. After vigorous vortexing for 10 to 30 s, samples were centrifuged at 17,000 × g for 10 min at 4°C. The supernatant was transferred to a new tube and dried completely on a SpeedVac.

Dried metabolite extracts were reconstituted in 100 µL of 50% (v/v) acetonitrile/water, vortexed, and centrifuged at 17,000 × g for 10 min. A 70-µL aliquot of the supernatant was transferred to HPLC vials, and 10 µL was injected per analysis. Chromatographic separation was performed on a Thermo Scientific Vanquish UHPLC system equipped with a SeQuant ZIC-pHILIC polymeric column (2.1 × 150 mm, 5 μm) maintained at 35 °C, using a flow rate of 150 µL min^-1^. The mobile phase consisted of 20 mM ammonium carbonate, pH 9.7 (A) and 100% acetonitrile (B), and samples were resolved with the following gradient program: 20% A to 80% A over 20 min, followed by a return to 20% A from 20 to 20.5 min, and re-equilibration at 20% A from 20.5 to 28 min. The effluent was analyzed by a Q-Exactive mass spectrometer operating in polarity-switching mode with the following settings: spray voltage, 3.2 kV; sheath gas, 40; auxiliary gas, 15; sweep gas, 1; capillary temperature, 275°C; auxiliary gas heater, 350°C. Full-scan MS1 spectra were acquired from *m/z* 70–1,000 at a resolution of 70,000, with an automatic gain control (AGC) target of 1 × 10^-6^ and a maximum injection time of 250 ms. Data-dependent MS2 scans were collected for the top three most abundant ions using a normalized collision energy of 35.

Raw files were centroided and converted to mzXML format using msconvert (ProteoWizard). Data processing was performed in MZmine 2: chromatograms were built using the ADAP chromatogram builder, peaks were detected with the ADAP wavelets algorithm, aligned across samples using the RANSAC aligner, gap-filled, and annotated by matching to an in-house MS1/retention-time database (mass tolerance ±15 ppm; RT tolerance ± 0.5 min). Identifications and peak boundaries were manually verified. Peak areas were integrated, normalized to both the internal standard (trifluoromethanesulfonate) and the DNA content of the extracted cells, and exported for statistical analysis.

DNA content was quantified using a fluorescence-based assay. Following methanol evaporation, samples were resuspended in Solution 1 (100 mM NaCl, 20 mM Tris-HCl pH 7.4, 5 mM EDTA, 0.1% SDS, 500 µg mL^-1^ Proteinase K), vortexed, and digested overnight at 37 °C with shaking. The next day, two volumes of Solution 2 (5 µg mL^-1^ Hoechst 33342) were added. After a 30-min incubation at 37 °C in the dark, 100 µL of each sample and a standard curve of UltraPure Salmon Sperm DNA (Invitrogen; prepared in Solution 1, ranging from 12.5 to 800 µg mL^-1^) were transferred to a black-walled, clear-bottom 96-well plate. Fluorescence was measured (excitation: 355 nm; emission: 465 nm) with a SpectraMax M2e microplate reader, and sample DNA concentration was determined by interpolation from the standard curve. Raw and processed metabolomics data are deposited into Metabolomics Workbench^103^ (Study ID: ST005225).

### ALDH expression and purification

*ALDH9A1* cDNA encoding the canonical isoform was obtained from SinoBiological (Cat: HG20431-UT) and cloned into a pET-15b plasmid vector. The pET-15b-ALDH9A1 plasmid was transformed into competent BL21 *E. coli*, and the cells were cultured in Luria–Bertani (LB) medium containing 100 µg mL^−1^ ampicillin at 37 °C and treated with 1 mM isopropyl β-D-1-thiogalactopyranoside (IPTG) overnight at 18 °C to induce protein expression. The bacterial cells were then harvested and lysed with BugBuster Master Mix buffer (Millipore Sigma) containing a protease inhibitor cocktail (cOmplete, Roche). The lysate was centrifuged (16,000 g at 4 °C for 30 min), and the soluble fraction was mixed with 2 mL of Ni-NTA Superflow resin. After rocking at 4 °C for 1 h, the mixture was used to create a gravity column, which was washed with wash buffer 1 (20 mM sodium phosphate, 500 mM NaCl, 1 mM DTT, 20 mM imidazole and 5% (v/v) glycerol, pH 7.4) and wash buffer 2 (20 mM sodium phosphate, 500 mM NaCl, 1 mM DTT, 40 mM imidazole and 5% glycerol, pH 7.4). The protein was then recovered from the column with elution buffer (20 mM sodium phosphate, 500 mM NaCl, 1 mM DTT, 400 mM imidazole and 5% glycerol, pH 7.4), concentrated using a 50-kDa Amicon Ultra unit, and dissolved in storage buffer (500 mM NaCl, 20 mM sodium phosphate, 1 mM DTT, pH 7.4). The final protein concentration was determined with a Pierce 660-nm Protein Assay Kit (ThermoFisher, 22662), and the samples were analyzed by SDS–PAGE and stored at −80 °C. The cloning, expression, and purification of the human recombinant isozymes ALDH1A1, ALDH1A2, ALDH1A3, ALDH1B1, ALDH2, ALDH3A1, ALDH3A2, ALDH4A1, ALDH5A1, and ALDH7A1 were performed as previously described^23^.

### ALDH enzyme kinetics assays

To evaluate ALDH enzymatic activities *in vitro*, reaction mixtures consisting of assay buffer (100 mM sodium phosphate, 1 mM MgCl_2_, 0.5 mM DTT, 0.005% (v/v) Tween 20, pH 8.0), 1 mM NAD^+^ and the indicated ALDH enzyme were prepared in white opaque 96-well assay plates (100 µL per well, flat-bottom and non-treated; Corning Costar). Corresponding aldehyde substrate (acetaldehyde, 4-HNE, BAL, ABAL or GBBAL) was then added to each well to achieve the desired concentrations and the resulting enzymatic activity was determined by changes in NADH levels over the course of 10 min. NADH fluorescence was monitored with a SpectraMax M2e microplate reader (340-nm excitation and 460-nm emission; Molecular Devices) operated by SoftMax Pro software.

ALDH enzymatic activity against all-*trans* retinal was assessed *in vitro* by preparing 100-µL reaction mixtures in white opaque, flat-bottom 96-well plates (Corning Costar). Each well contained assay buffer (100 mM sodium phosphate, 1 mM MgCl_2_, 0.5 mM DTT, 0.005% (v/v) Tween 20, pH 8.0), 1 mM NAD⁺, 0.05 mM resazurin (Fisher Scientific, R12204), 1 U/mL diaphorase (Worthington Biochemical, LS004330), and the specified ALDH enzyme. Reactions were initiated by adding all-*trans* retinal to the desired final concentrations, and the resulting enzymatic activity determined by the conversion of resazurin to resorufin over the course of 10 min. Resofurin fluorescence was monitored using the SpectraMax M2e microplate reader (560-nm excitation and 590-nm emission) operated by SoftMax Pro software.

### Mouse liver mitochondrial lysate assays

Animal work performed for this study complies with the Stanford University Institutional Animal Care and Use Committee regulations (Stanford IACUC protocol #14145). Mitochondria were isolated from fresh mouse liver tissue through differential centrifugation, using organs isolated from C57BL mice and a double knock-in transgenic line inserting a frameshift that prematurely terminates the encoded protein (*ALDH1B1^G193fs^* mice^46^). Briefly, approximately 500 mg of tissue was minced and homogenized with an electric homogenizer on ice in 1 mL of cold homogenization buffer (210 mM mannitol, 70 mM sucrose, 50 mM MOPS, 1 mM EDTA, pH 7.4) supplemented with the Roche protease inhibitor cocktail. The homogenate was centrifuged at 700 g for 2 min at 4 °C to pellet nuclei and cellular debris, and the resulting supernatant was centrifuged at 7,000 g for 10 min at 4 °C to pellet the crude mitochondrial fraction. This pellet was washed twice with 1 mL of cold homogenization buffer followed by centrifugation at 7,000 g for 5 min at 4 °C. To lyse the mitochondria, the final pellet was resuspended in 0.5 mL of lysis buffer (100 mM Tris-HCl pH 8.0, 10 mM DTT, 20% (v/v) ethylene glycol, 0.5% (v/v) Triton X-100) and vortexed briefly. The lysate was clarified by ultracentrifugation at 100,000 g for 30 min at 4°C. The final supernatant, representing the soluble mitochondrial fraction, was then collected. The protein concentration was determined with the Pierce 660-nm Protein Assay Kit and stored at -80 °C for subsequent analysis. ALDH activity measurements were conducted with the protocol described for ALDH enzyme kinetics assays, using 10 µg total mitochondrial protein in 100 µL of assay buffer.

### Western blot analysis

Mitochondrial lysates from mouse liver were prepared as described above, and lysates from cultured cells were prepared as previously reported^23^. Protein samples were mixed with 6× SDS–PAGE loading buffer (240 mM Tris-HCl, 6% SDS, 0.3 M DTT, 30% (v/v) glycerol and 0.018% bromophenol blue, pH 6.8) to afford a final protein concentration of 2.0 µg µL^−1^. 25 µL of each sample was then loaded and resolved on 4–15% Criterion TGX Stain-Fre^e^ Protein Gel (Bio-Rad) at 130 V for 1.5 h at room temperature. Total protein was detected with a Gel Doc XR imaging system (Bio-Rad) before the gel was transferred to PVDF membranes using the Trans-Blot Turbo Transfer System (Bio-Rad). The membranes were blocked with 5% non-fat dry milk in 1× Tris-buffered saline containing 0.1% (v/v) Tween 20 (TBST) for 1 h at room temperature. The blots were then probed with primary antibody in the same blocking buffer overnight at 4 °C. The membranes were washed 4 × 5 min with TBST and incubated with horseradish peroxidase-conjugated secondary antibody in blocking buffer for 2 h at room temperature. After the membranes were washed 4 × 5 min TBST, the secondary antibodies were detected by chemiluminescence using SuperSignal West Dura Extended Duration Substrate (Thermo Fisher Scientific) and the Gel Doc XR imaging system. Antibodies used in these experiments include rabbit polyclonal anti-ALDH1B1(1:500 dilution; Proteintech, Cat No.22220-1-AP), rabbit polyclonal anti-ALDH9A1 (1:500 dilution, Proteintech, Cat No. 26621-1-AP), rabbit polyclonal anti-SLC25A20 (CACT, 1:500 dilution; Invitrogen, Cat No. PA5-53508) and horseradish peroxidase-conjugated donkey polyclonal anti-rabbit IgG (1:1,000 dilution; GE Healthcare, NA934-1ML).

### ALDEFLUOR assays

ALDH1B1-overexpressing *ALDH1A3^−/−^* A375 cells were generated as previously described^33^ and cultured in a 10-cm dish for 24 h, and then trypsinized and resuspended in ALDEFLUOR assay buffer (STEMCELL Technologies) at a concentration of 4 × 10^5^ cells/mL. A 0.5-mL aliquot of this cell suspension was treated with individual compounds for 15 min at 37 °C under a 5% CO_2_ atmosphere. The cells were next treated with 2.5 µL of ALDEFLUOR reagent and incubated for an additional 30 min at 37 °C under 5% CO_2_. The cells were then collected by centrifugation (450 g for 5 min at 4 °C) and resuspended in 200 µL of assay buffer. One drop of DAPI viability dye (ThermoFisher, R37606) was added to each sample prior to flow cytometry analysis.

### Immunofluorescence imaging

To assess the subcellular localization of exogenously expressed ALDHs, *ALDH1B1^−/−^* SW480 cells overexpressing FLAG-tagged ALDH1B1, ALDH2, and ALDH9A1 were seeded at a density of 1.2 × 10^5^ cells/well into a 24-well plate containing poly-D-lysine-coated glass coverslips. After 24 h, cells were incubated with MitoTracker Deep Red (500 nM; Invitrogen, M22426) for 30 min at 37 °C to label mitochondria, then fixed with 4% paraformaldehyde in PBS for 10 min at room temperature. Following permeabilization with 0.3% (w/v) Triton X-100 in PBS, cells were incubated with anti-FLAG polyclonal antibodies (1:200; Proteintech, 20543-1-AP) overnight at 4 °C. After washing, cells were incubated with Alexa Fluor 488-conjugated donkey anti-rabbit IgG (1:200; ThermoFisher, A-21206) for 1 h at room temperature in the dark. Coverslips were mounted with Antifade Mounting Medium containing DAPI (Vector Laboratories, H-1800-2) and imaged using a Zeiss LSM 800 confocal microscope with ZenBlue software.

### CACT transport assays

The coding sequence of CACT was amplified by PCR using cDNA from wild-type SW480 cells and cloned into the pMW172 vector (Addgene, 89925) via Gibson Assembly (NEB, E2611S). The resulting pMW172-CACT plasmid was transformed into C41(DE3) competent *E. coli* (Sigma, CMC0017), and an overnight culture (5 mL) was inoculated into 1 L of LB medium containing ampicillin (100 μg mL^-1^) and grown at 37 °C. At OD_600_ = 0.55–0.65, protein expression was induced with 0.5 mM IPTG, and then cultures were incubated at 37 °C for 24 h. The bacterial cells were harvested by centrifugation (2,800 g at 4 °C for 20 min).

Isolation of inclusion bodies was performed with the following modifications to the published procedure^75^. Cell pellets were resuspended in 20 mL of lysis buffer (10 mM Tris-HCl, 0.1 mM EDTA, 1 mM PMSF, Roche protease inhibitor cocktail, pH 8.0) and lysed by sonication at 4 °C (Qsonica Q500, 6-mm tip, 40% amplitude, 3 s on/9 s off cycles for 20 min). Cell debris was removed by centrifugation (27,000 g at 4 °C for 15 min), and the pellets were resuspended in 4 mL of resuspension buffer (10 mM Tris-HCl, 0.1 mM EDTA, protease inhibitor cocktail, pH 8.0). The sucrose step-gradient centrifugation was performed as previously described, but at 131,000 g at 4 °C for 2 h in a Beckman Coulter Optima XPN-90 ultracentrifuge with the ramp and deceleration set to 5/10. The CACT-containing inclusion body pellets were stored at −20 °C for subsequent refolding steps.

Each inclusion-body pellet was solubilized in 800 µL solubilization buffer (2% sarkosyl, 30 mM PIPES, pH 7.0) and incubated end-over-end at 4 °C for 20 min. Insoluble material was removed by ultracentrifugation (124,000 g at 4 °C for 30 min). An aliquot (594 µL) of the supernatant was supplemented with 6 µL of 100 mM DTE (final concentration of 1 mM) and incubated at 4 °C for 60 min. The sample was then diluted with 120 µL of buffer (10 mM PIPES, 500 mM NaCl, pH 7.0) and applied to a Sephadex G-75 column (0.6 × 9 cm) pre-equilibrated with 10 mM PIPES, 100 mM NaCl, pH 7.0. The first 2 mL of eluate was discarded, and the subsequent 600 µL was collected for downstream reconstitution.

L-α-phosphatidylcholine (chicken egg; Avanti; 60% purity) was supplied in chloroform and 2 mL was transferred to a 50-mL round-bottom flask. Chloroform was removed by rotary evaporation under vacuum at room temperature (∼45 min for 2 mL), and the resulting lipid film was resuspended in buffer (20 mM NaCl, 10 mM PIPES, 1 mM EDTA, pH 7.0) to a final concentration of 100 mg/mL. The flask was sealed with Parafilm, incubated at room temperature for at least 1 h, and then vortexed in 30 s intervals (2 min total) and mixed with a glass Pasteur pipette until homogeneous. The flask was sealed with a septum cap, flushed with N₂ for at least 5 min, and sonicated in a bath sonicator for 60 min. Bath temperature was monitored and maintained below 30 °C by adding ice as needed.

CACT inclusion bodies were reconstituted into proteoliposomes and assayed for exchange activity using [^3^H]-carnitine, with the following modifications to published procedures.^75,104^ Proteoliposome mixtures were assembled on ice in a final volume of 700 µL containing: 60 µg protein (typically 20–30 µL of Sephadex G-75 eluate; protein quantified with a Pierce 660-nm Protein Assay Kit, Thermo Fisher), 100 µL of 10% L-α-phosphatidylcholine liposomes, 0.7 mg cardiolipin, 1% (v/v) Triton X-100, 10 mM PIPES (pH 7.0), and 13 mM internal substrate. Amberlite XAD-2 columns (0.5 g resin packed in 0.6-cm diameter glass pipettes to 3.5-cm bed height) were used for cyclic detergent removal as previously described^104^. Transport assays with the CACT-containing proteoliposomes were conducted using the inhibitor-stop method^105^, initiating the assay by adding 10 µL of 1.1 mM [^3^H]-carnitine (0.1 µCi per sample) and quenching CACT activity by adding 10 µL of 60 mM NEM dissolved in transport buffer (60 mM NaCl, 10 mM PIPES, pH 7.0). Each sample collected for final analysis was mixed with 10 mL Ultima Gold XR scintillation cocktail and quantified using a Hidex SL600 scintillation counter.

### FAO assays

FAO activity was measured in both adherent cells and spheroids. For adherent cells, SW480 or HepG2 cells were seeded in six-well plates (3.0 × 10⁵ cells/well) and, at ∼60% confluence, treated with DMSO, 2 µM IGUANA-5, or 2 µM IGUANA-C for 24 or 48 h. For spheroids, SW480 cells (5 × 10³ cells/chamber) were seeded in removable chamber slides (ThermoFisher, #178599) pre-coated with 50 µL Matrigel and cultured in spheroid media for 4 days prior to a 24 h treatment with the same compounds. Following these incubations, both models were treated for 2 h with DMSO, 2 µM IGUANA-5, 2 µM IGUANA-C, or 10 µM etomoxir in their respective culture media, washed twice with HEPES-buffered saline (HBS), and then incubated with 10 µM FAOBlue reagent (DiagnoCine, #FNK-FDV-0033) in HBS for 1 h. After a final HBS wash, adherent cells were collected for flow cytometry analysis, while spheroids were mounted under coverslips and imaged using a Zeiss LSM 800 confocal microscope with ZenBlue software.

### Flow cytometry

Flow cytometry was performed for the ALDEFLUOR and FAO assays. DAPI (Thermo Fisher, R37606) and TO-PRO-3 (Thermo Fisher, R37170) were used as viability dyes for the ALDEFLUOR and FAO assays, respectively. Cellular fluorescence was measured using the following settings: DAPI (405-nm excitation, 450/50-nm emission), FAO signal (405-nm excitation, 450/50-nm emission), ALDEFLUOR signal (488-nm excitation, 525/50-nm emission), and TO-PRO-3 (640-nm excitation, 670/30-nm emission). Cells were gated by FSC-A vs SSC-A to exclude debris, FSC-A vs FSC-H to identify singlets, and DAPI or TO-PRO-3 vs SSC-A to select live cells. For each sample, 10,000 events were acquired, and ALDEFLUOR and FAO signals were quantified using live singlets only.

### RNA-seq analyses

Parental SW480 cells were cultured under adherent and three-dimensional spheroid conditions in three biological replicates for each group. For adherent culture, cells were seeded in six-well plates at a density of 2.0 × 10^5^ cells/well in 3.0 mL of culture medium. For spheroid culture, wells were first pre-coated with 250 µL of Matrigel (Corning). Cells were then seeded at a higher density of 4.0 × 10^5^ cells/well in 3.0 mL of medium to promote spheroid formation. After 48 h of incubation, cells were harvested for total RNA extraction. For spheroids, the Matrigel matrix was first dissolved by adding 1.5 mL of Cell Recovery Solution (Corning) and incubating on ice for 15 min. Intact spheroids were then collected by centrifugation at 800 × g at 4 °C for 5 min.

Total RNA from both adherent cells and pelleted spheroids was isolated using the Monarch Total RNA Miniprep Kit (New England Biolabs) according to the manufacturer’s protocol. The total mRNA was processed and analyzed by Novogene using the NovaSeq 6000 platform. For each sample, approximately 40 million paired-end reads were obtained, and raw FASTQ data were processed through fastp. Paired-end clean reads were aligned to the GRCh38 reference genome using the Spliced Transcripts Alignment to a Reference (STAR) software, and FeatureCounts was used to count the read numbers mapped of each gene. Differential expression analysis was performed using the DESeq2 R package. The resulting *P* values were adjusted using the Benjamini-Hochberg approach for controlling the false discovery rate. Raw and processed RNA-seq data are deposited into the Gene Expression Omnibus database (GEO accession GSE345612).

## Statistics and reproducibility

All data were initially recorded and processed by Microsoft Excel. Graphing and statistical analysis were performed using GraphPad Prism Version 11.0.0 (93). No data points were excluded from all analysis, and data were presented as the mean ± s.e.m., with replicate numbers indicated in the figure legends. Biochemical assays (Figs. 1E, 3A-E, and 3G-I and Supplementary Figs. 2A-B, 3, 4, and 6B) were performed at least three times, cell-based assays (Figs. 1B-D, 5A-C, 5G-H, 6B-G and Supplementary Figs. 1A-D, 6C-D and 7A) were performed at least two times, and the CACT assay with three internal substrates (Fig. 7C) was performed twice. Student’s *t*-tests (two-tailed and unpaired) were used to compare results and calculate *P* values.

## ACKNOWLEDGEMENTS

We thank E. Patton for providing the *ALDH1A3^−/−^* A375 cells, Y. Chen for assistance with mitochondria extractions from mouse liver, C. Indiveri for valuable discussions about the CACT assay, and S. Demuro and M. Rabbani for their critical reading of the manuscript and helpful comments.

## CONTRIBUTIONS

Z.F. and J.K.C. conceived the study. Z.F., T.E.B., S.L.C., A.M.K., L.L., H.R.C, and J.K.C. designed the experiments and interpreted the data. Z.F., S.L.C. and N.M. performed the metabolomics profiling studies. Z.F. conducted the ALDEFLUOR, western blot, FAO, immunofluorescence imaging, ALDH rescue, ATP-based cell viability, and RNA-seq studies. T.E.B. and A.M.K. synthesized and characterized GBBAL, and A.K.T. synthesized IGUANA-5. L.L. conducted imaging-based cell viability assays with the HT-1080^N^ cell line. Z.F. and T.E.B. expressed and purified ALDH proteins and conducted *in vitro* enzymatic assays. Z.F., C-H.C. and T.S. extracted mitochondrial proteins from mouse liver, and Z.F. conducted following enzymatic assays with these samples. T.E.B., A.M.K. and Z.F. conducted the CACT assays. Z.F. and J.K.C. wrote the manuscript with input and editing from all authors. S.J.D, D.M.R., H.R.C., and J.K.C. acquired funding for the project.

