## Supplementary Information for "ALDH1B1 promotes mitochondrial γ-butyrobetaine biosynthesis and fatty acid oxidation in colorectal cancer"

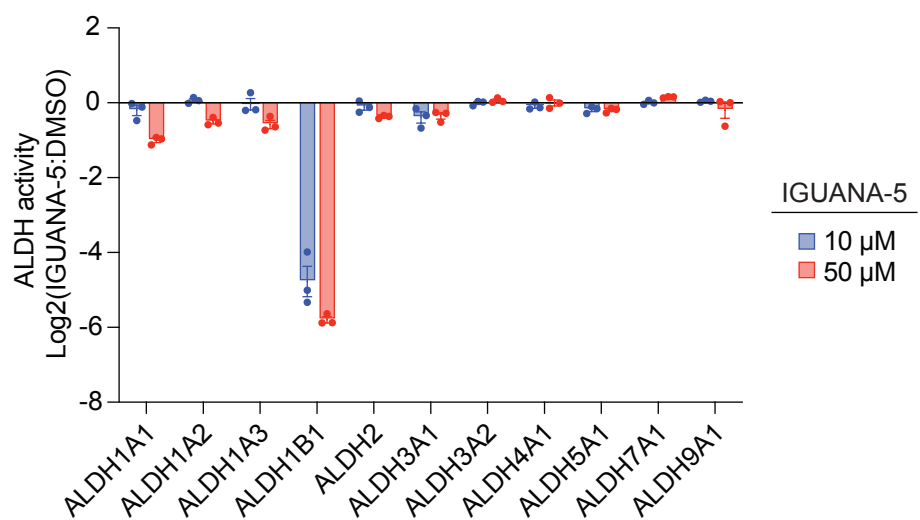

**Supplementary Figure 1. IGUANA-5 is an ALDH1B1-selective inhibitor.** Activities of IGUANA-5 against selected ALDH isoforms. Assays were conducted with an ALDH monomer concentration of 100 nM and 1 mM acetaldehyde substrate, and data are the average of three biological replicates ± s.e.m.

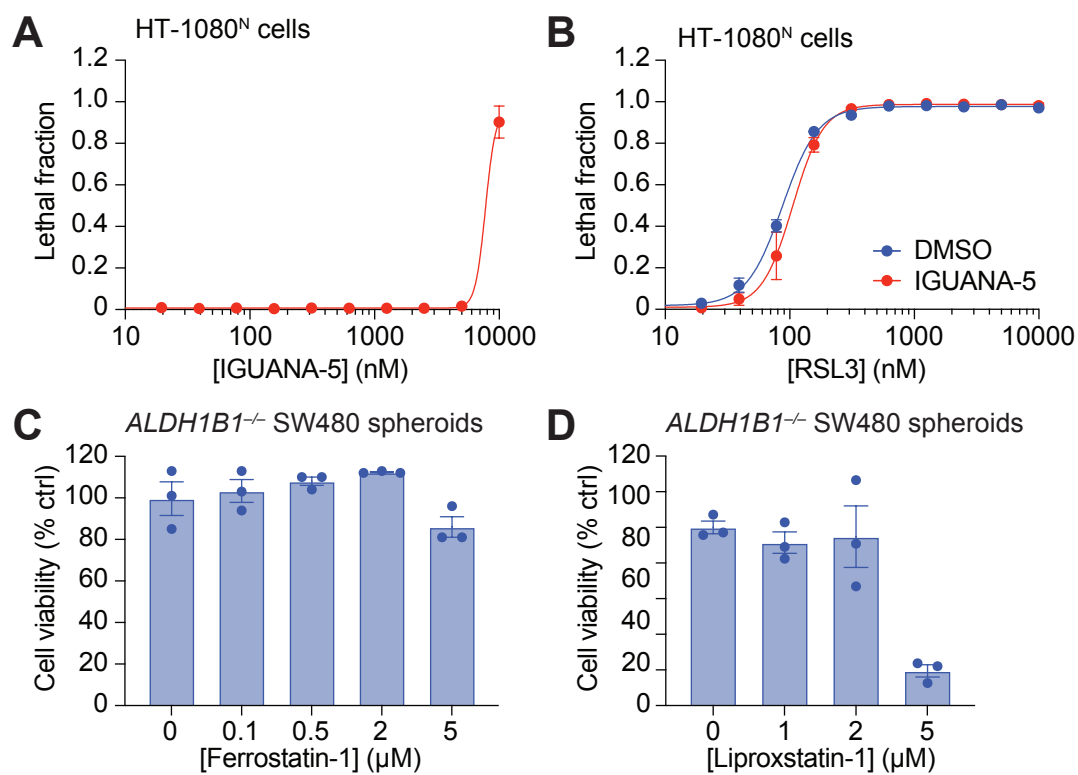

**Supplementary Figure 2. ALDH1B1 inhibition does not promote ferroptosis.** (A) Dose-dependent effects of IGUANA-5 on the viability of HT-1080<sup>N</sup> cells cultured as 2D monolayers. (B) Dose-dependent effects of RSL3 on 2D HT-1080<sup>N</sup> cultures the presence or absence of IGUANA-5. The cells were pre-treated with DMSO or 2 μM IGUANA-5 for 24 h and then co-treated with RSL3 for additional 48 h. The lethal fraction scores in (A-B) were calculated by quantifying the cell populations with SYTOX Green-positive staining. (C-D) Dose-dependent effects of the ferroptosis inhibitors ferrostatin-1 (C) and liproxstatin-1 (D) on *ALDH1B1*<sup>-/-</sup> SW480 spheroids. Data are average of two biological replicates ± s.e.m. in (A-B) and three biological replicates ± s.e.m. in (C-D).

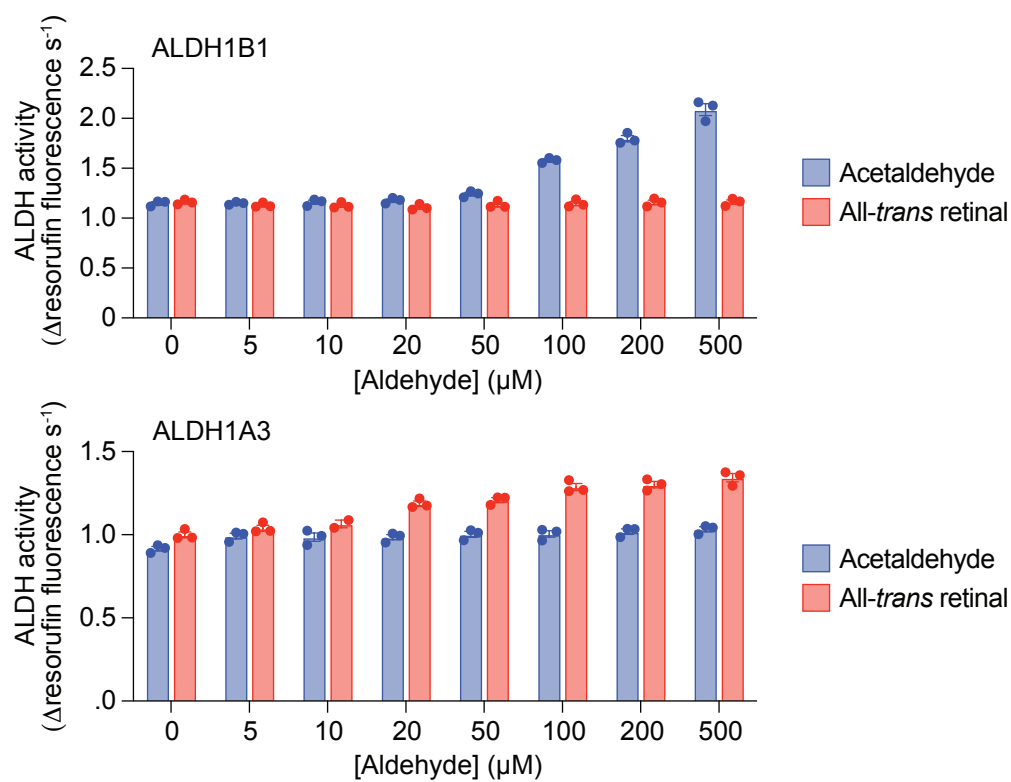

**Supplementary Figure 3. ALDH1B1 does not efficiently oxidize all-*trans* retinal.**

Oxidizing activity of recombinant ALDH1B1 against acetaldehyde and all-*trans* retinal as substrates, as determined with an NADH-diaphorase coupled assay. The oxidizing activity of recombinant ALDH1A3 is shown for comparison. Assays were conducted with an ALDH monomer concentration of 100 nM, and data are the average of three biological replicates ± s.e.m. Background aldehyde-independent increases in resorufin levels are due to reduction of the diaphorase substrate resazurin by dithiothreitol, an assay buffer additive that is required to maintain ALDH1B1 activity.

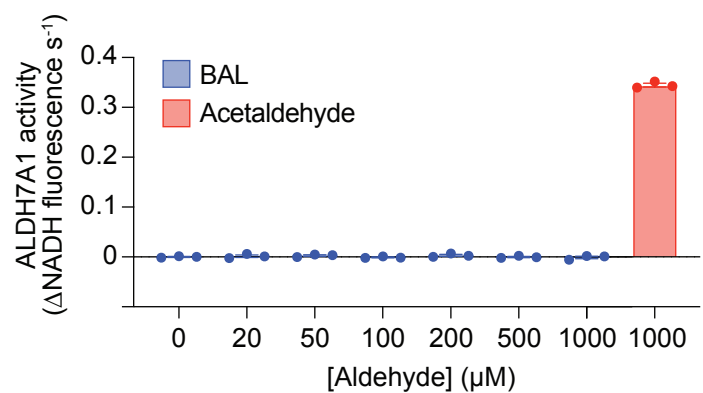

**Supplementary Figure 4. ALDH7A1 does not efficiently oxidize BAL *in vitro*.** Oxidizing activity of recombinant ALDH7A1 against BAL and acetaldehyde substrates, as determined by in NADH production. Assays were conducted with an ALDH7A1 monomer concentration of 100 nM, and data are the average of three biological replicates ± s.e.m.

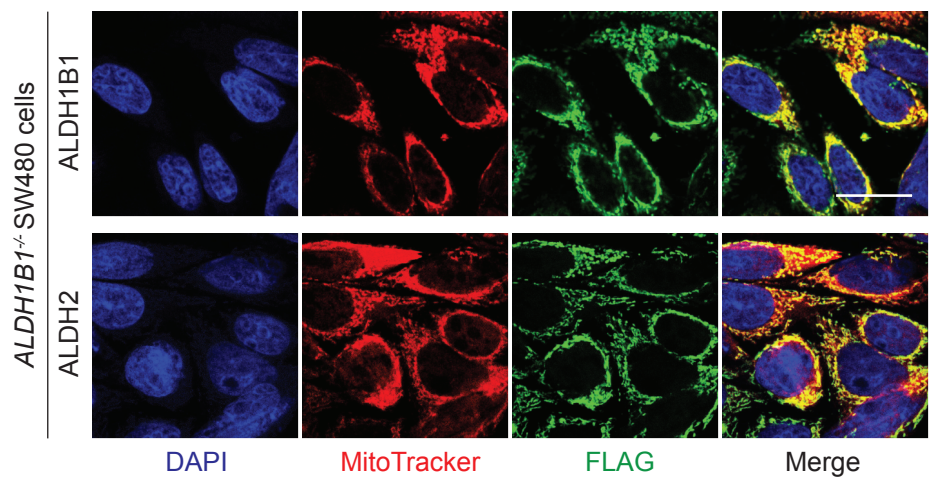

**Supplementary Figure 5. Subcellular localization of exogenously overexpressed ALDH isoforms.** Immunofluorescence images of *ALDH1B1*<sup>-/-</sup> SW480 cells transduced with lentivirus encoding ALDH1B1 or ALDH2 with a C-terminal FLAG tag. Mitochondria were labeled with MitoTracker, and nuclei were counterstained with DAPI. Scale bar: 20 μm.

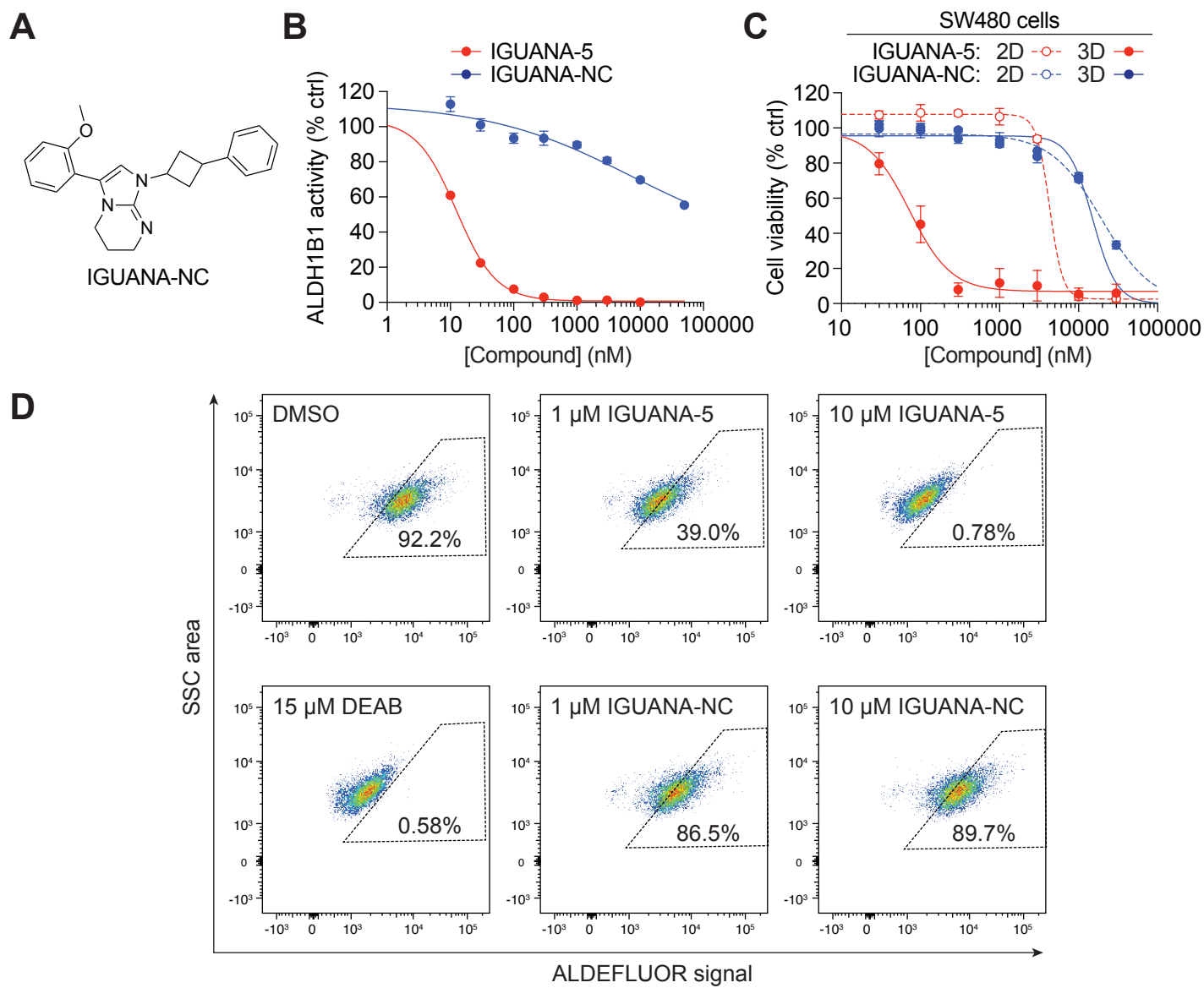

**Supplementary Figure 6. IGUANA-NC is an inactive analog of guanidinyldene ALDH1B1 inhibitors.** (A) Chemical structure of IGUANA-NC. (B) Dose-dependent effects of IGUANA-5 and IGUANA-NC against recombinant ALDH1B1 enzyme *in vitro*. Assays were conducted with an ALDH1B1 monomer concentration of 100 nM and 1 mM acetaldehyde substrate. (C) Dose-dependent effects of IGUANA-5 and IGUANA-NC on adherent and spheroid cultures of SW480 cells. (D) FACS plots of ALDH1B1-overexpressing *ALDH1A3*<sup>-/-</sup> A375 cells that were incubated with the indicated compounds and then treated with ALDEFLUOR reagent. 10,000 cells were analyzed for each condition, and ALDEFLUOR-positive populations were gated as shown by the dashed boxes. Data are the average of three biological replicates ± s.e.m. in (B-C).

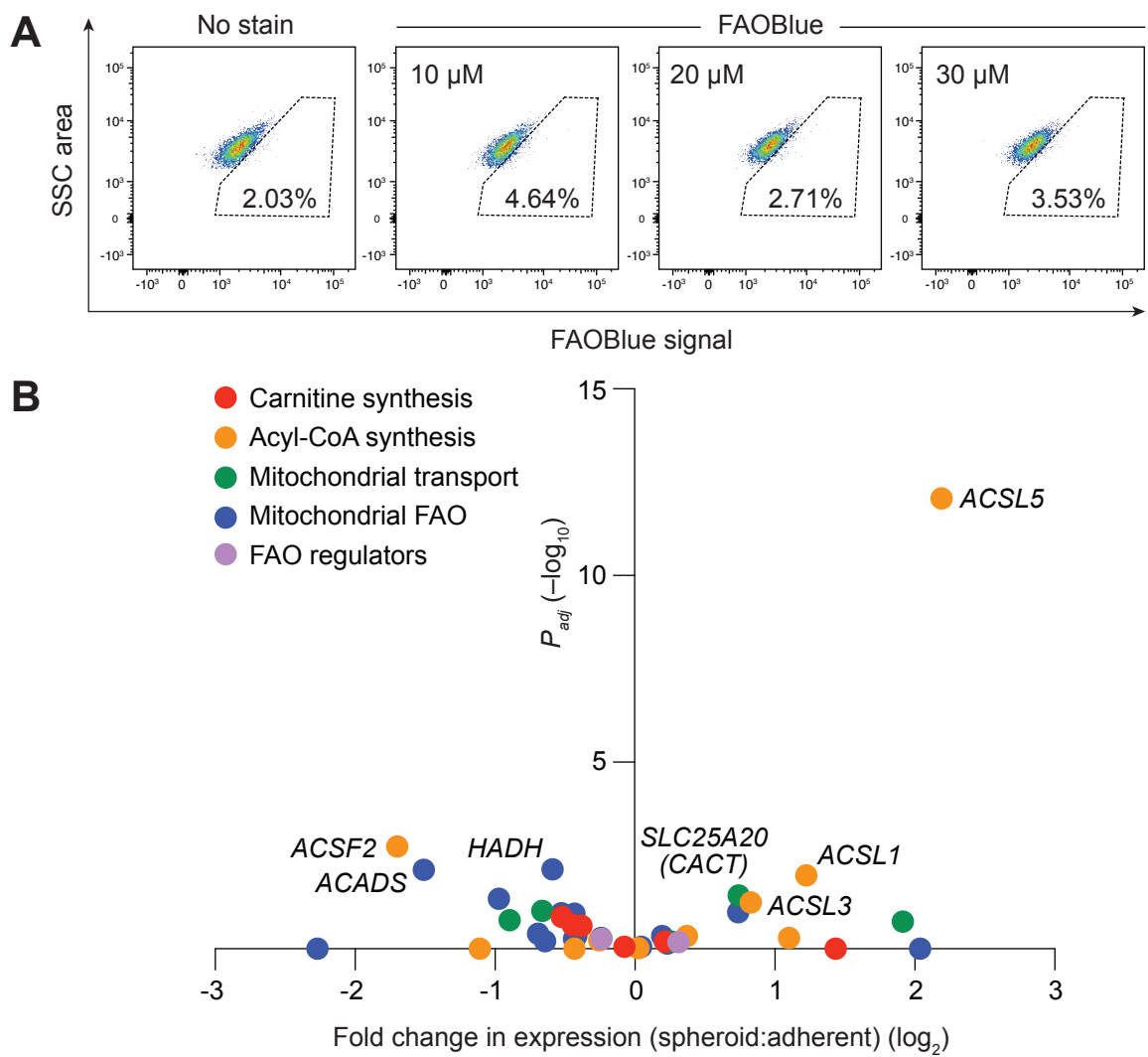

**Supplementary Figure 7. Adherent cultures of SW480 cells have minimal FAO activity.** (A) Representative FACS plots of SW480 cells cultured as 2D monolayers and treated with various concentrations of FAOBlue. 10,000 cells were analyzed for each condition, and FAOBlue-positive populations were gated as shown by the dashed boxes. (B) Differential expression of carnitine- and mitochondrial FAO-related genes in spheroid versus adherent cultures of SW480 cells, as determined by whole-transcriptome RNA sequencing. Selected genes with the largest transcriptional changes are annotated.

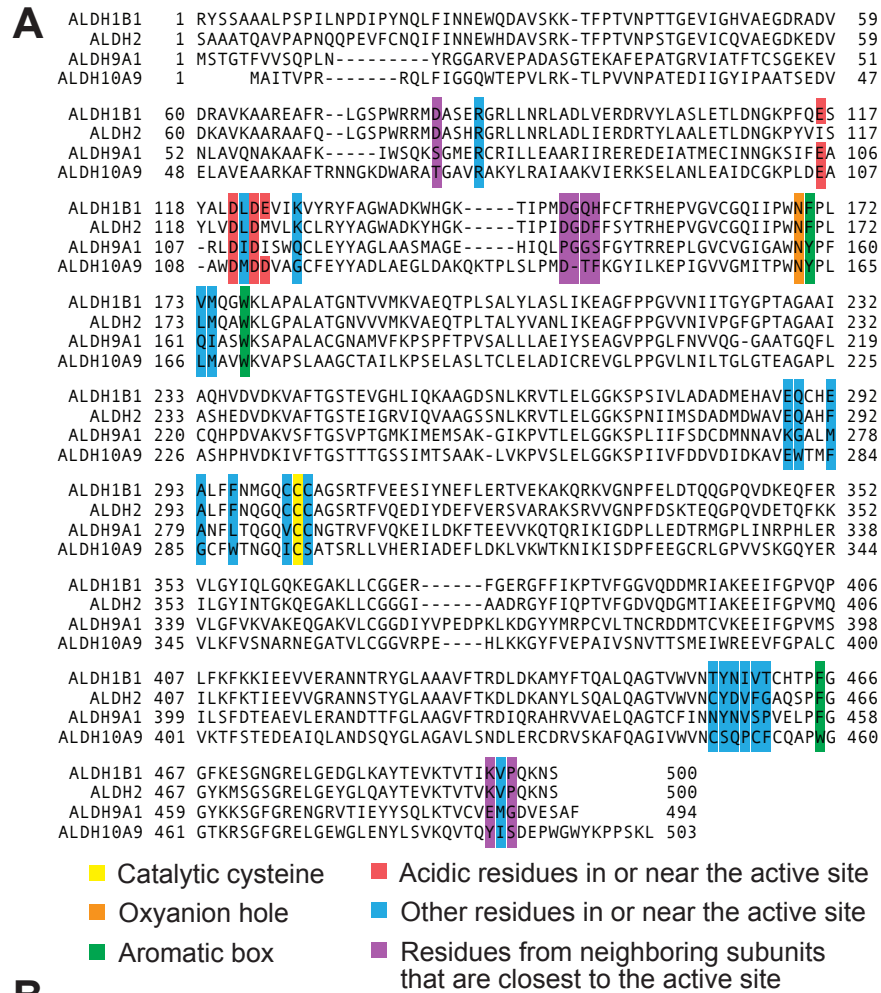

**B**

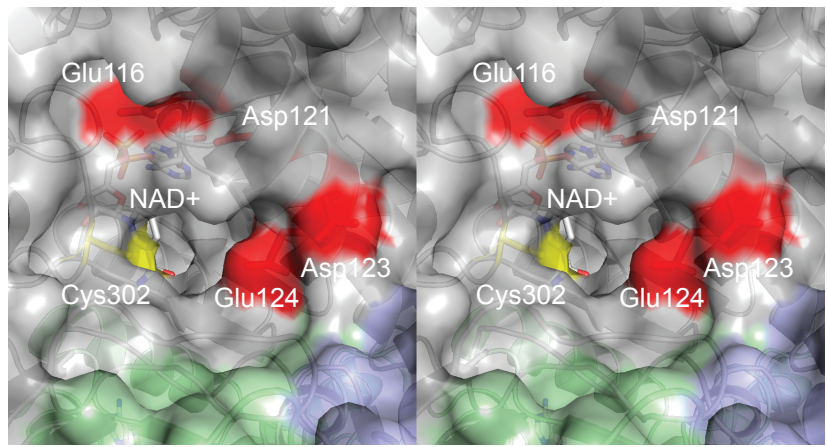

**Supplementary Figure 8. Conserved and divergent structural elements in the ALDH family.** (A) Sequence alignment of ALDH1B1, its closest homolog ALDH2, ALDH9A1, and an ALDH10 family member from *Arabidopsis*. (B) Wall-eyed stereoview of the ALDH1B1 active site (PDB: 7MJC) shown as a ribbon and transparent surface model, with visible tetramer subunits colored grey, blue, and green. A unique cluster of acidic residues (red), the catalytic cysteine (yellow), and NAD<sup>+</sup> are shown as stick models.
